# Modeling Translational Riboswitches: The impact of SAM concentration on the folding of the SAM-II riboswitch

**DOI:** 10.1101/2024.05.09.593440

**Authors:** Osama Alaidi

## Abstract

Several mechanistic (thermodynamic) models have been developed for the folding of SAM-II riboswitch as a function of SAM and magnesium ions concentrations. For each model, the parameters have been determined from experimental (apparent) binding data, based on the underlying assumptions of the model. The predicted titration curves computed from the different models were calculated and compared with actual experimental observation of the fraction of the RNA forming a pseudoknot at specific concentrations of the ligands. Strikingly, only one of the six models correctly predicts the experimental findings, unveiling the dominant mechanism of the riboswitch function. More interestingly, the latter mechanism is found to be the most efficient compared to the other possible mechanisms. The study sheds light on the cognate ligand conformational capture mechanism of the SAM-II riboswitch in the presence of specific concentrations of magnesium ions. The presented mathematical and thermodynamic framework, as well as the inferred equilibrium constants, provide foundations for making accurate quantitative prediction of the SAM-II riboswitch ensemble populations as a function of SAM and magnesium concentrations. The mechanistic linked equilibria model can be generalized to describe other thermodynamically driven riboswitches and hence facilitate identifying RNA intermediates that can be leveraged for small molecule drug design.

**Graphical Abstract:** 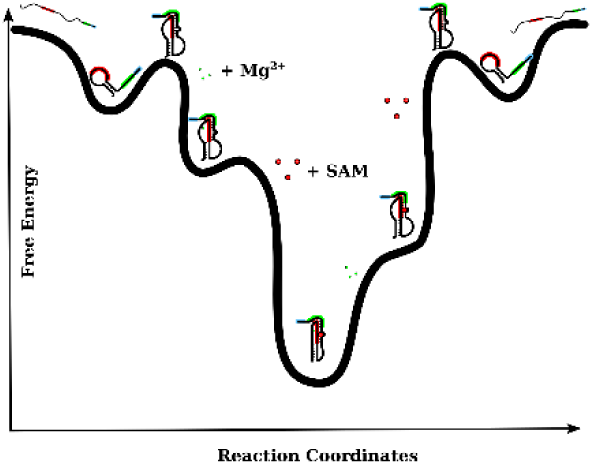

## 1 INTRODUCTION

Riboswitches are RNA sensor elements, primarily present in bacterial mRNA, that aid in gene regulation of small molecule metabolism on both the transcriptional and translational levels (1). To date, more than fifty classes of riboswitches have been confirmed, and thousands of novel riboswitches are predicted in bacterial genomes (2). The SAM-II riboswitch is a class of riboswitches that binds to the metabolite; SAM (S-Adenosyl-Methionine), leading to termination of ribosomal translation of the downstream genes that are involved in the metabolism of SAM (3, 4). The mechanism by which the RNA switches its conformation from the ON to the OFF state, in the presence of SAM is yet to be fully understood (5–10). Understanding such a mechanism is crucial to be able to control the function of the riboswitch by means of SAM or other designed small molecules (11, 12). It is likely that riboswitches share common functional mechanisms. Consequently, structural and mechanistic insights from one riboswitch may be observed in other classes of riboswitches, and hence the importance of mechanistic studies on simple model riboswitches such as the SAM-II riboswitch from the Sargasso Sea metagenome which controls the mRNA translation of the *metX* gene (13, 3–7, 14, 8, 9).

In an earlier study, the impact of magnesium ions on the folding of the SAM-II riboswitch was investigated (10). In this manuscript, mechanistic models, by which the riboswitch RNA conformational changes may take place in response to its cognate ligand (SAM), are presented and applied to predict titration curves for the SAM-II riboswitch. The models build on the general framework used in our previous work (10), and generally combines RNA structure prediction along with linked equilibria approach to enable the prediction of the fraction of riboswitch bound and/or folded at equilibrium, at various ligand concentrations. By deriving and applying models that carry various hypotheses, different possible mechanisms by which the ligand may induce its effect were examined. Comparing the results from these models indicated that one of the mechanisms is more plausible than the others and hence likely to be the dominant mechanism.

## 2 METHODS: Developing ligand-induced conformational shift models for RNA

### 2.1 Predicting ligand concentration-dependent riboswitch folding (titration curves): The overall workflow

Titration curves provide a means to compare model predictions with experimental findings. Doubtlessly, the accuracy of the predictions made, based on a certain model, reflects the accuracy of the underlying assumptions of the model. In this study, mechanistic linked equilibria models were developed and used to examine the effect of SAM on the SAM-II riboswitch population in the presence of Mg^2+^ ions. The derived models that were used for the calculations in this study are tabulated in **Table 1**, along with the major assumptions made in each model. Previously studied models describing the impact of magnesium ions on the riboswitch (designated as *Mg^2+^ binding models*) are also listed in the table for clarity. The models are schematically illustrated in **Figure 1**.

**Figure 1.**
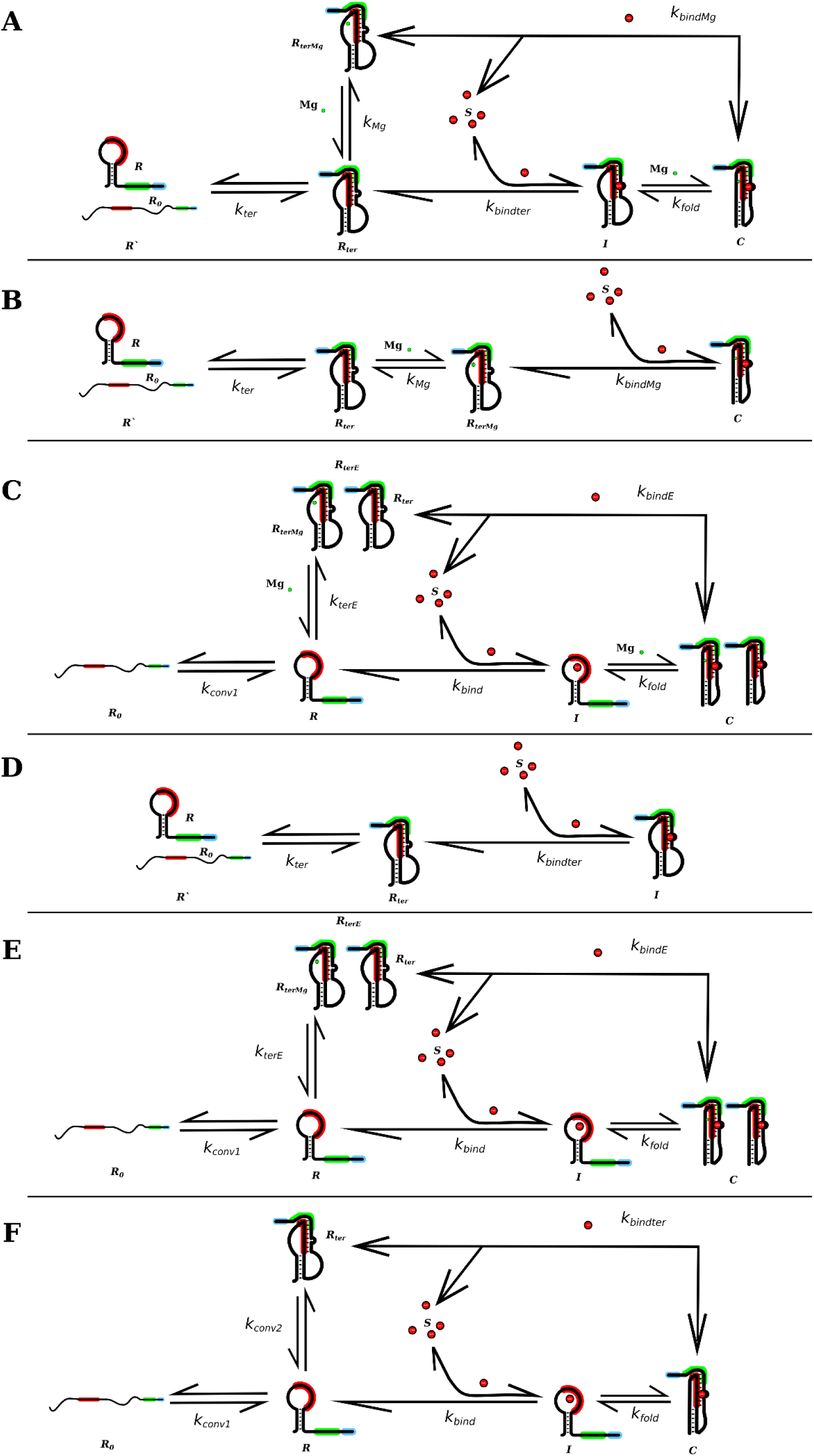
Schematic representations of the models described in this study (Models 1-6). The figure panels illustrate graphical reaction schemes for the proposed models designated as *Model 1-6.* The related Thermodynamic cycles for *Models 1, 3, 5* and *6* are shown, respectively, in Figure 2A-2D. (A) *Model 1.* The model assumes a conformational capture mechanism that is conditioned by the presence of a pseudoknot that is stabilized by the binding of SAM. The latter binding can occur with or without the prior binding of Mg^2+^. That is, in this mechanism, SAM binds to two different forms of the partially formed pseudoknot structures (i.e. two intermediates *R*_*ter*_ and *R*_*terMg*_), hence induces the complete RNA folding and SAM trapping. As shown in the results this model is most likely to reflect the functional mechanism of the SAM-II riboswitch. (B) *Model 2*. The model assumes that SAM binds via a conformational capture mechanism. The occurrence of the latter mechanism is conditioned by both the presence of pseudoknot and the binding of Mg^2+^. (C) *Model 3*. The model considers the differential binding between the conformers with and without the pseudoknot as well as the effect of Mg^2+^ concentration. However, any RNA that contains the specific secondary structure is assumed to bind with SAM. (D) *Model 4*. The model assumes a conformational capture mechanism based on the presence of a pseudoknot and assumes that the effect of Mg^2+^ concentration is negligible. Hence, the model describes a Mg^2+^ independent, single pathway for SAM binding. (E) *Model 5*. The model considers the differential binding between the conformers with and without the pseudoknot but assumes that the effect of Mg^2+^ concentration is negligible. (F) *Model 6*. The model assumes that SAM can bind to all RNA that has the minimal secondary structure. In other words, SAM binding occurs in the presence or absence of tertiary interactions, with no effect of Mg^2+^.

**Table 1.** Summary of the models constructed to describe the mechanism of the SAM-II riboswitch. The models vary in their underlying assumptions regarding the role of tertiary interactions in the conformational capture mechanism, the role of alternative SAM-binding pathways and whether the model considers the presence and concentration effects of Mg^2+^. The *Model 0* and the *Mg^2+^ Model,* are listed here for clarity (and have been described elsewhere) (10) and they represent, respectively, the description of the RNA in the absence of both ligands (Mg^2+^ and SAM) and a model that describe the impact of Mg^2+^ on the RNA, in the absence of SAM. In the table above, the - and + signs are used to indicate whether the feature or assumption is considered by in the model. *SAM Models 1* to *6* are pictorially illustrated in panels **A**-**F** of **Figures 1**, respectively.

| Model name | SAM binding | Presence of $Mg^{2+}$ | Effect of $Mg^{2+}$ concentration | Tertiary interactions play a role in SAM conformational selection | Alternative SAM binding pathways | Reference |
| --- | --- | --- | --- | --- | --- | --- |
| *Model 0* | - | - | - | Not applicable | Not applicable | Reference (10). (also reviewed in the Supplementary methods). |
| *Mg<sup>2+</sup> Model* | - | + | + | Not applicable | Not applicable | Reference (10) |
| *Model 1* | + | + | + | + | + | This manuscript |
| *Model 2* | + | + | + | + | - | This manuscript |
| *Model 3* | + | + | + | - | + | Derived in reference (10) |
| *Model 4* | + | - | - | + | - | This manuscript |
| *Model 5* | + | + | - | - | + | This manuscript |
| *Model 6* | + | - | - | - | + | This manuscript |

The presented six SAM-binding models in this study assume distinct mechanisms, and differ from each other based on whether if: (1) the presence of Mg^2+^ is essential for SAM binding, (2) the Mg^2+^ concentration influence the fraction of the finally folded state, (3) tertiary interactions are essential for the conformational capture mechanism to take place, (4) more than one mechanistic pathway may play a role in the apparent binding. Each of the developed models therefore examines a certain set of assumptions or a proposed plausible mechanism. The model that correctly predicts an experimental observable, is that whose underlying assumptions are likely to be correct. Hence, the comparison of the modeling results with experimental values can provide an indication of which model is likely to be valid.

The usual workflow in using such mathematical models was followed here. First, the analytical/mathematical model was constructed based on a plausible mechanism. Obviously, any mechanism will intrinsically have certain assumptions. Second, the model parameters were determined based on a few measured apparent equilibrium constants obtained from the literature. Third, a titration curve was computed based on each model and were compared to an experimentally observed/measured values that, in this case, were not used to infer the model parameters. The model which gave values that are most consistent with experimental observations, was considered the one that is most likely to occur, and hence effectively represents the mechanism of the riboswitch function.

The presented models (**Table 1** and **Figure 1)** were named according to the presence and absence of Mg^2+^ and SAM in the model. For example, *Model 0* (**Figure S1**) is a model that simply describes the riboswitch folding in the absence of both Mg^2+^ and SAM, whereas *Mg^2+^ Model* is a model that describes the riboswitch folding in the presence of Mg^2+^ and in the absence of SAM. The latter models are mentioned here for clarity and have been described in our previous study (10). The SAM-binding models developed in this work (*Models 1* to *6*) depict six distinct mechanistic models that can describe the riboswitch folding in the presence of SAM, and in which the effect of Mg^2+^ may (either implicitly or explicitly) or may not be taken into account, depending on the specific scheme and the underlaying assumptions of the model.

The SAM-binding models (**Figures 1**, panels **A-F)** are described in detail in the following sections of the manuscript. Moreover, as elaborated in the following sections, the equilibrium constants for each model are inferred based on the apparent experimentally determined binding affinities, and when necessary, from the cognate thermodynamic cycles shown in **Figures 2**, panels **A-D**.

**Figure 2.**
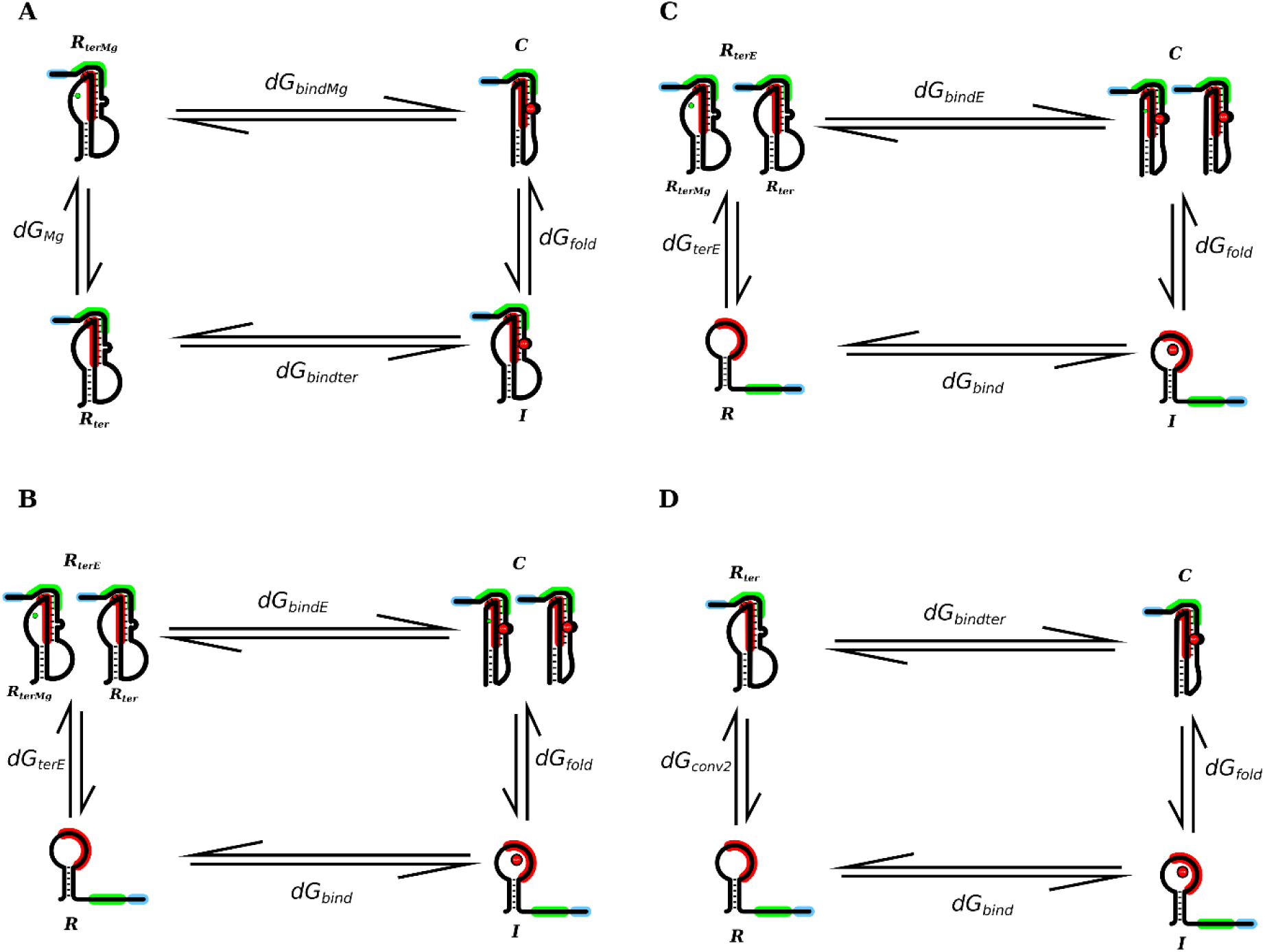
Cognate thermodynamic cycles. The figure illustrates the cognate thermodynamic cycles that were used to facilitate the determination model parameters based on the measured apparent binding constants. The cycles shown in panels **A**-**D** are for *Models 1,3,5* and *6*, respectively.

### 2.2 Definitions of the RNA species (states) and parameters (equilibrium constants) used in the models

The following definitions or nomenclature of the RNA species and model parameters (equilibrium constants) are, for consistency, used in all models, though not all the models necessarily have all these species and parameters. The following nomenclature is therefore used throughout this manuscript.

***i- Total and free concentrations.*** Like in the Mg^2+^ binding model described in our previous work (10), in all models presented in this study, the total RNA, Mg^2+^ ions and SAM concentrations are designated as *R*_*tot*_, *Mg*_*tot*_ and *S*_*tot*_, respectively. Whereas the free (unbound) Mg^2+^ ions and SAM are designated as *Mg* and *S*, respectively.

***ii- RNA species (states) concentrations.*** In general, RNA species with a pseudoknot formed are considered to be in the OFF state (inaccessible ribosome binding site) whereas RNA species without a pseudoknot formed are in the ON state (accessible ribosome binding site). The RNA conformers in the models are defined as follows.

*R*_0_is the concentration of the unbound *binding incompetent* RNA (i.e. RNA conformers in which the SAM binding contacts are not accessible). Such RNA states/species do not show the specific RNA secondary structure that is known to bind to SAM, nor are they bound with any ligands. *R* is the concentration of the unbound intermediate. The latter class of conformers has the minimal secondary structure required for SAM binding (i.e. which allows the SAM binding contacts to be accessible) but does not have a pseudoknot. *R*′ is the RNA concentration without a pseudoknot. Hence, this class of RNA is composed of more than one state, namely, *R*_0_ and *R* (i.e. *R*’ = *R*_0_ + *R*). *R*_*ter*_ is the concentration of the riboswitch RNA with a nucleated pseudoknot but not bound to Mg^2+^ nor SAM.

*R*_*terMg*_is a state structurally similar *R*_*ter*_ except that it is bound with Mg^2+^. *R*_*terE*_ , is the concentration of RNA with a nucleated pseudoknot that effectively enters the subsequent reaction, regardless of its binding with Mg^2+^. This class of RNA is effectively a mixture of states, i.e. *R*_*terE*_ = *R*_*ter*_ + *R*_*terMg*_. *I* is the concentration of the SAM bound intermediate (SAM bound riboswitch with or without the pseudoknot, depending on the model). *C* is the concentration of the SAM bound *final and most stable state of the riboswitch* (SAM bound riboswitch with the pseudoknot fully formed). Depending on the model, this class of conformers may or may not be bound to Mg^2+^ or may represent a mixture of both. It is important to note that only species *I* and *C* are bound to SAM.

***iii- The equilibrium constants used in the Models (model parameters).*** The equilibrium constants for the RNA conversions are defined as follows. *K*_*conv*1_ is the equilibrium constant for the conversion from *R*_0_ to *R* and *K*_*conv*2_ is the equilibrium constant for the conversion from *R* to *R*_*ter*_. To reduce the number of parameters in the tested models as much as possible, and to ease the incorporation of experimental data, the following apparent equilibrium constants are defined. *K*_*ter*_ is the apparent equilibrium constant for the conversion from *R*′to *R*_*ter*_ and *K*_*terE*_ is the apparent equilibrium constant for the conversion from *R* to *R*_*terE*_. Hence, both *K*_*ter*_ and *K*_*terE*_ are apparent equilibrium constants.

As described above, each of *R*′ and *R*_*terE*_are composed of more than one state. The equilibrium constants for RNA binding, are defined as: *K*_*Mg*_ is the equilibrium binding association constant of Mg^2+^ to species *R*_*ter*_, *K*_*bind*_is the equilibrium binding association constant of SAM to species *R*, *K*_*bindE*_ is the equilibrium binding association constant of SAM to species *R*_*terE*_, *K*_*bindter*_ is the equilibrium binding association constant of SAM to species *R*_*ter*_, *K*_*bindMg*_ is the equilibrium binding association constant of SAM to species *R*_*terMg*_, and *K*_*fold*_ is the equilibrium association constant for the folding of the intermediate *I* to form *C*.

In the absence of any ligands (**Figure S1**, *Model 0*), the equilibrium constants for RNA conversion can be simply obtained from RNA folding probabilities, as follows.

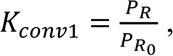, represents the ratio of probabilities of conformers *R* and *R*_0_ , in the *absence* of SAM and absence of tertiary interactions. Hence, the calculation can be performed exclusively based on secondary structure energetics.

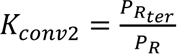, represents the ratio of probabilities of *R*_ter_ and *R*, in the *absence* of SAM. The probabilities of these populations do not rely on any ligands, and the value for the equilibrium constant was obtained from the previously reported two-step Mg^2+^ binding model (10).

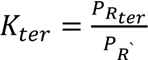, represents the ratio of probabilities of *R* and *R*′, in the *absence* of SAM. The probabilities of these populations also do not rely on any ligands, and the value for the equilibrium constant has also been obtained from the previously reported two-step Mg^2+^ binding model (10).

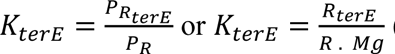, (depending on the model, reflecting if the model considers or ignores the effect of magnesium), represents the ratio of probabilities of *R*_*terE*_and *R*, in the *absence* of SAM. The ratio between these two populations relies on the concentration of Mg^2+^ ions. The probabilities and equilibrium constant can also be obtained from a two-step Mg^2+^ binding model (10).

For each model, the binding and folding parameters are determined from the apparent experimental binding constants reported in the literature, with the help of thermodynamic cycles if needed. The latter equilibrium constants may include, *K*_*bind*_, *K*_*bindter*_, *K*_*bindMg*_, *K*_*bindE*_ and *K*_*fold*_. Thereafter, a quantitative prediction of the concentration of the bound and folded (i.e. with the pseudoknot formed) riboswitch (i.e. species *I* and/or *C*, depending on the model) is obtained as a function of the equilibrium constants and the total concentrations of RNA and ligands (Mg^2+^ and SAM). To enable this, it was essential to establish links between the latter equilibrium constants and the literature experimental (apparent) binding constants.

### 2.3 Defining the experimental equilibrium binding constants

The constants, *Ka*_*ITC*_and *Ka*_*FS*_denotes two experimental (apparent) binding association constants previously reported in the literature that are frequently used in this work, that are based on Isothermal calorimetry (ITC) (4) and Fluorescence spectroscopy (FS) (6), respectively. Whereas the former defines binding in the usual ratio between the bound and free RNA in the presence of the ligand, the second monitors the changes in the RNA folding. More specifically, the latter method monitors the changes (increase or decrease) in fluorescence intensity from a 2-Aminopurine inserted at a specific site (as an indicator for the increase or decrease of a state), that arise from local interactions (many of which are tertiary interactions) associated with RNA folding, as a function of the ligand concentration (15–17). Effectively, such technique monitors the fraction change in the emitted fluorescent signal, at a specific wavelength range, as the cognate ligand is added. In their study on the SAM-II riboswitch, Haller *et al.* monitored the increase in the fluorescent signal associated with the increase of SAM concentration at several bases that undergo changes associated with the pseudoknot formation (6). Since the equilibrium constant from such method is based on monitoring the change in the fraction of RNA that gives off a signal at the specific wavelength range, and that the ensemble contains a fraction of RNA that already has a pseudoknot prior to adding SAM as well as a fraction that remains unfolded, then the measured constant represent, the ratio between the concentration of the folded & bound RNA relative to the concentration of folded & unbound RNA, taking the ligand concentration into account. Thus, the reaction scheme and equilibrium constant, respectively, can be written as: 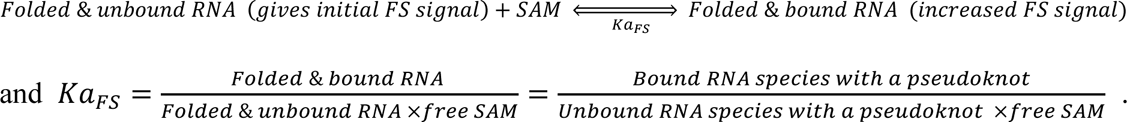 In the latter expression, the concentrations of the folded and bound RNA species will depend on the specific scheme for each model. As briefly discussed in the following section and fully described in the **Supplementary methods**, by specifying those RNA species for each model, links between the experimental measurements and the model were established, and the model parameters were inferred.

### 2.4 Computing model parameters and the fractions of individual RNA species

The parameters for each model are obtained based on the apparent experimental equilibrium constants as described in the *Determination of Model parameters* section for each model in the **Supplementary methods**. Given the equilibrium constants for each model, the concentrations of the final (folded) RNA species *C* (or *I* in *Model 4*) were computed (using Eq. 1.15, Eq. 2.6, Eq. 3.15, Eq. 4.4, Eq. 5.15, Eq. 6.15, in the **Supplementary methods** for *Models 1-6,* respectively) as a function of the concentration of model ligands. The expressions for computing the final/folded RNA state are also tabulated in **Table S2**. Further, concentrations/fractions of all stable RNA species in the models were calculated as a function of SAM and Mg^2+^ concentrations based on the definition of the equilibrium constants in each model (see model derivations in the **Supplementary methods**).

Finally, the *fraction bound* (corresponding to the total probability of bound RNA at a given SAM concentration (*S*_*tot*_)), as well as the *fraction folded*, the *fraction both bound* & *folded* and the *fraction binding competent*, can be computed the summation of respective fractions of appropriate species for each model as summarized in **Table S3**.

### 2.5 Models for quantitative prediction of RNA-ligand interactions for the SAM-II riboswitch

The following section describes the models derived and used in this study.

#### Model 1: A ligand-binding model for RNA that considers the effect of both Mg^2+^ ions and tertiary interactions

The model, designated as *Model 1*, is schematically illustrated in **Figure 1A**. In this model, the cognate ligand (SAM) binds to all molecules with a nucleated pseudoknot, whether bound with Mg^2+^ ions or not. Hence, this model considers the effect of both Mg^2+^ concentration and tertiary interactions, and the SAM-induced folding can be achieved through either magnesium-dependent or a magnesium-independent pathway.

#### Model 2: A ligand-binding model for RNA that considers the effect of both Mg^2+^ ions and tertiary interactions in which the primary ligand can only bind if Mg^2+^ is bound

This model, designated as *Model 2*, is schematically illustrated in **Figure 1B**. In this model, the cognate ligand (SAM) binds exclusively to molecules with a nucleated pseudoknot and that are bound with Mg^2+^ ions. Hence this model considers the effect of both Mg^2+^ and tertiary interactions, however, assumes that the SAM-induced folding of the RNA is only achievable through a single magnesium-dependent pathway. Given the equilibrium constants (*K*_*Mg*_and *K*_*ter*_) and after computing *K*_*bindMg*_(from Eq. 8), the concentration and fraction of *C* were computed (Eq. 2.6 in the **Supplementary methods**, **Table S2**) at different concentrations of Mg^2+^ and SAM.

#### Model 3: A ligand-binding model for RNA that ignores the effect of tertiary interactions in the conformational capture mechanism but considers the effect of Mg^2+^ ions

This model, designated as *Model 3*, is schematically illustrated in **Figure 1C**. In this model, unlike the previous two models, the cognate ligand conformational selection relies on the formation of secondary structure only, and not on tertiary interactions. However, the model considers the effect of Mg^2+^ concentration explicitly. It is worth noting that this model is similar to *Model 5* except that here the explicit Mg^2+^ concentration is taken into account.

#### Model 4: A ligand-binding model for RNA that considers the effect of tertiary interactions but ignores the role of Mg^2+^

This model, designated as *Model 4*, is schematically illustrated in **Figure 1D**. This model assumes a conformational selection mechanism in which SAM binds only with RNA that possess a nucleate pseudoknot, hence, the effect of tertiary interactions is considered in this model. However, the binding of a single ligand, SAM, is taken into account and the effect of Mg^2+^ binding is assumed to be negligible. In effect, the model represents a simple classical two step conformational capture mechanism in the absence of Mg^2+^. The equations for this model are derived and described in detail in reference (10), except that the ligand used here is SAM. Therefore, the model assumes that only a single magnesium-independent pathway, for the riboswitch binding and folding, is possible. From the scheme and the definitions of *K*_*bindter*_and the definition of *K*_*FS*_ used in the above models, it can be seen that *K*_*bindter*_ ≈ *K*_*FS*_.

#### Model 5: A ligand-binding model for RNA that assumes ligand recognition is not influenced by tertiary interactions and ignores the concentration effect of Mg^2+^

This model, designated as *Model 5,* is schematically illustrated in **Figure 1E**. In this model, the concentration effect of Mg^2+^ is assumed to be negligible and the observed effect of Mg^2+^ can be implicitly taken into account using an apparent equilibrium constant (*K*_*bindE*_). Moreover, like *Model 3*, it is assumed that all RNA molecules with plausible secondary structures can bind with SAM, regardless of the presence of tertiary interactions. Nonetheless, the model takes into account that the binding affinities of the conformers that have tertiary interactions may differ from those that lack such interactions.

Figure 1E schematically illustrates *Model 5* for SAM binding and folding of the SAM-II riboswitch. This model assumes that SAM binding is not influenced by the concentration of Mg^2+^ ions and assumes that the formation of the correct secondary structure is enough to form binding competent conformers. As described in section **2.2**, the equilibrium constants for the model shown in (Figure 1E) are defined as follows. *K*_*conv*1_, *K*_*terE*_, *K*_*bind*_, *K*_*bindE*_ and *K*_*fold*_ are the equilibrium constants for the conversion between (*R*_0_and *R*), between (*R* and *R*_*terE*_), binding of the intermediates, binding of pseudoknot containing structures and folding of the bound intermediates (formation of pseudoknot in the presence of the ligand), respectively.

#### Model 6: A ligand-binding model for RNA that assuming that SAM recognition is not influenced by tertiary interactions and occurs in the absence of Mg^2+^ ions

This model, designated as *Model 6*, is schematically illustrated in Figure 1F. In this model, the impact of Mg^2+^ is assumed negligible, and the magnesium bound species are not represented in the model. Despite that the latter assumption contradicts previously reported experimental data, it is still instructive to derive and study this model for the purpose of comparisons with other models, as well as, for confirming the most probable mechanism by excluding other possibilities. As in the previous model, here, the tertiary interactions are assumed to have no major role in SAM recognition. In other words, it is assumed that all RNA molecules with plausible secondary structures (and not necessarily any specific tertiary interactions) can bind with SAM in the absence of magnesium.

## 3 RESULTS

### 3.1 Determining model parameters (equilibrium constants) enables computing titration curves for the individual RNA species of the different models

To be able to predict the corresponding titration curves for each model, the model parameters were determined (as described in the **Supplementary methods** section). One key concept we have used in this study is expressing the apparent (and experimentally measured) binding in terms of the binding constants that are specific to each model. Hence, the observed *fraction bound* can incorporate different species or conformers that vary according to the model. This allowed us to establish links between the apparent experimental measured quantities with the different conformers proposed in the model, enabling the computation of the model equilibrium constants. If the model correctly reflects the underlaying mechanism, then it should be able to predict other independent experimental observations. For instance, here we used binding data, to compute the parameters for all models. As a result, the fraction bound is the same across all models, and even certain individual species may be the same, but the fraction folded is the key difference. As a test, we then contrasted the predicted fraction folded against the experimentally observed fraction reported in the literature that formed a pseudoknot at a specific concentration of the SAM and Mg^2+^. Further, we have taken into account the methods by which binding was measured. Specifically, whereas ITC measurements evaluate the *Kd* based on the bound and free molecules, several previously reported FS-based apparent *Kd* for riboswitches (6, 18) were based on the local or global RNA structural changes, observed as an increase in the fluorescence signal change, upon ligand binding. Hence, the interpretation of the apparent *Kd* by each technique is different.

*Model 1* depends on five parameters (equilibrium constants): *K*_*ter*_, *K*_*Mg*_, *K*_*bindter*_, *K*_*bindMg*_ and *K*_*fold*_(Figure 1A). As illustrated in Figure 1A and in **Figure S1**, *K*_*ter*_is the apparent association equilibrium constant for the formation of the pseudoknot in the absence of Mg^2+^ ions. The latter constant depends on both *K*_*conv*2_ and *K*_*conv*1_ (**Figure S1**). In the absence of tertiary interactions, SAM and Mg^2+^ ions, the only two accessible states of the system are *R*_0_ and *R*. The equilibrium ratio between the two spices is controlled by the constant *K*_*conv*1_. Hence, the latter constant can be determined from secondary structure calculations, by calculating the ratio between the probability of conformers that form a closed loop (i.e. the P1 helix is formed, denoted as species *R*) and those that do not have the P1 helix (denoted as species *R*_0_). The calculations were performed using the method previously reported by Alaidi *et al*. (19, 10). Further, using the Mg^2+^ model described in our previous study (10), *K*_*ter*_ and *K*_*Mg*_ were determined. Thereafter, using one of the experimentally reported constants *Kd*_*ITC*_ (4) and *Kd*_*FS*_(6), along with the thermodynamic cycle shown in Figure 2A, the remaining parameters (*K*_*bindter*_, *K*_*bindMg*_ and *K*_*fold*_) were determined as described in detail in the methods section. The determined equilibrium constants and/or corresponding free energies for this model are listed in **Table 2**.

**Table 2.** *Model 1* parameters. The table lists the parameters computed for *Model 1*. Despite that SAM binding energies, in the presence or absence of Mg^2+^ (*dG*_*bindMg*_ and *dG*_*bindter*_, respectively) clearly play the major role in stabilizing the final folded conformer, a significant energetic contribution to the stability comes from both the Mg^2+^ binding and the Mg^2+^ mediated tertiary interactions that leads to pseudoknot stabilization or zipping of the binding pocket (*dG*_*Mg*_ and *dG*_*fold*_, respectively). It should be noted that *K*_*ter*_ can be expressed in terms of *K*_*conv*1_and *K*_*conv*2_as described in detail in the *Mg Model* (10). The free energies were calculated from the corresponding equilibrium constants, such that Δ*G*_*i*_ = *RT* ln(*Kd*_*i*_) = − *RT* ln(*Ka*_*i*_), at room temperature (298.15 Kelvin).

| *Model 1* |  |
| --- | --- |
| Constant | Value |
| Given $K_{Mg}$ from *Mg Model* (10) | 878 |
| Given $K_{ter}$ based on *Mg Model* (10) | 0.0819 |
| Given $Kd_{ITC}$ from experiment (4) | 6.70E-07 |
| **Parameters in the presence of $Mg^{2+}$ (kcal/mol):** |  |
| $dG_{Mg}$ | -4.016 |
| $dG_{ter}$ | 1.483 |
| $dG_{fold}$ | -3.696 |
| $dG_{bindter}$ | -9.608 |
| $dG_{bindMg}$ | -9.289 |
| $dG_{FS}$ (calculated from model) | -9.425 |

The determination of the model parameters for *Models 2* and *Model 4* (Figures 1B and **1D**), however, is much simpler and more straightforward compared to the other models. The two models generally resemble the 2 or 3 step equilibrium models (10) and their parameters can be directly determined without the need to construct a thermodynamic cycle (as described in the **Methods** section).

On the contrary, as in *Model 1*, determining the parameters for *Models 3, 5* and *6* (Figures 1C, **1E** and **1F**) are not as straightforward, due to the differential binding affinities between the RNA species. This is because the latter three models assume that tertiary interactions influence the degree of binding with SAM. These models therefore resemble a conformational capture mechanism that is based on differential binding. This may occur when SAM is loosely selective for conformers with tertiary interactions (nucleated pseudoknot), though still all conformers with correct secondary structure bind with significant affinity to SAM. More specifically, the models account for the difference in binding affinity between the riboswitch with and without the nucleated pseudoknot as well as the difference in the folding in the presence of the ligand compared to its absence. In other words, the models consider that both riboswitches lacking the pseudoknot (with the binding contacts accessible and are in close proximity, enforced by a stable P1 stem and L1 loop) and riboswitch having a nucleated pseudoknot bind to SAM, with different affinities. This discrepancy in the binding, in turn, results in a difference in the energy of the formation of the pseudoknot (i.e. a difference in the folding energy) in the presence and absence of the ligand.

Given the limited available experimental data, as in *Model 1*, constructing thermodynamic cycles was needed to determine all the parameters for these models (Figures 2B, **2C** and **2D**). In *Models 3* and *5*, as in previous attempts (10), it was assumed that Mg^2+^ level is almost constant in the cell. On the one hand, in *Model 5* the system is modeled at a fixed Mg^2+^ concentration. On the other hand, *Model 3* makes it possible to study the system at different concentrations of Mg^2+^. In the latter models, it is assumed that the formation of the P1 helix allows SAM contacts as well as the residues that form the pseudoknot in the loop (L1) to be in close proximity to the ligand. Hence, all structures with the expected RNA secondary structure are assumed to be binding competent. Based on the previous experimental data and models (7, 8, 10), the equilibrium value *K*_*conv*2_at 2 mM Mg^2+^ can be obtained. Further, *K*_*bindE*_ or *K*_*bindter*_has been experimentally measured by FS experiments. From this value and from the experimentally measured overall binding *Kd*_*ITC*_the equilibrium constant *K*_*fold*_can be determined (see **Supplementary methods** for detailed derivation). Finally, *K*_*bind*_can be calculated with the aid of the corresponding thermodynamic cycles (Figures 2B, **2C** and **2D**). **Table S1** lists all parameters calculated and used for *Models 2-6*.

### 3.2 The comparison of predicted titration curves from the SAM binding models with experimental observations reveals that *Model 1* is the most probable model by which the SAM-II riboswitch function

For each model, the fractions of *individual RNA species*, as well as the *fraction bound*, the *fraction folded*, the *fraction bound & folded,* and the *fraction of binding competent RNA* have been determined as a function of SAM. To be able to determine which model truly represents how the SAM-II riboswitch works, we compared an experimentally observed fraction at certain conditions with the one predicted from various models. Specifically, it has been shown experimentally by single molecule FRET, that in the presence of 10 µM SAM (depicted by the vertical dashed line in Figure 3A) and 2 mM Mg^2+^, increased the folded RNA by ten-fold (6). Since in the absence of both ligands, the fraction of conformers that forms a pseudoknot is ∼ 9.3-9.5% (as reported in (8), and ∼ 9.34% after fitting to Mg^2+^ data as discussed (10)), then the expected fraction having a partially or fully formed pseudoknot (*fraction folded*) would be ∼ 93-95 % (specifically, 93.4%). Figure 3 shows the *fraction folded*, as well as the *fraction bound*, the *Bound and folded fraction* and the *fraction of binding competent RNA* as a function of increasing SAM concentration (panels **A-D,** respectively), at the experimental level of Mg^2+^. The only models that give a *fraction folded* that is very similar to the latter FRET experiment are *Model 1* and *Model 2,* predicting that 94.64% (and 94.62% when accounting for non-specific Mg^2+^) of the RNA population is folded (with a pseudoknot) at these experimental conditions, as opposed to 83.18%, 85.05%, 81.84% and 34.35% (depicted by the horizontal dashed line in Figure 3A) by *Models 3-6*, respectively.

**Figure 3.**
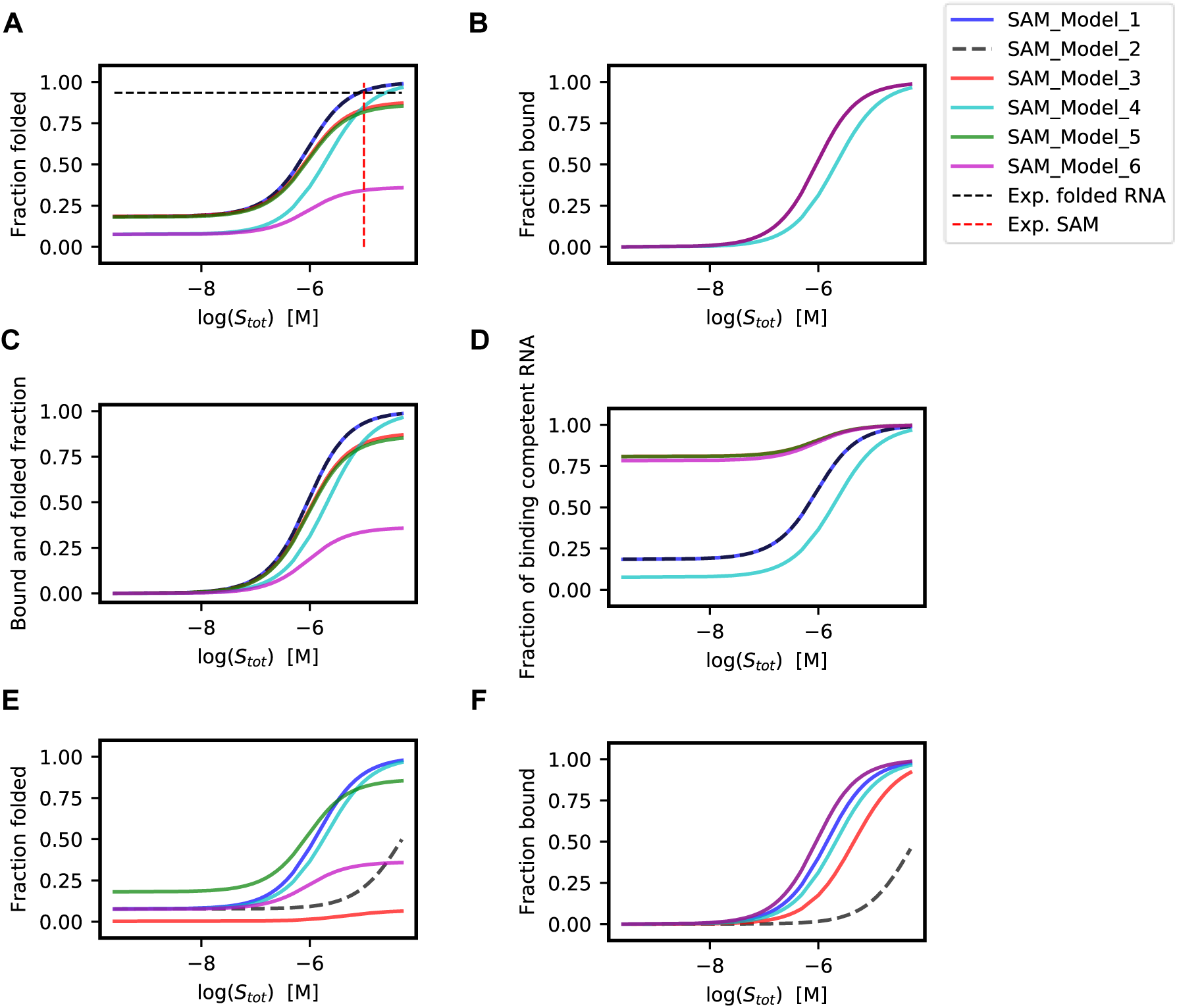
Titration curves for the SAM-II riboswitch assuming different models. The curves were simulated using a total SAM concentration of 10 µM. Panels **A** to **D** show the simulations results at a fixed total magnesium concentration of 2.0 mM using the different models. The panels show the computed *fraction folded*, *fraction bound*, *the bound and folded fraction* and the *fraction of binding competent RNA*, respectively, for all SAM *Models 1-6*. The dashed red and black lines (in panel **A**) depict the level of SAM and corresponding folded RNA in the previously reported single molecule FRET experiment (6) that is contrasted here against the predictions from the models. Panels **E** and **F**, show the computed *fraction folded* and *fraction bound*, respectively, at a much lower total magnesium concentration (20 µM) using the same models. All simulations were performed using an RNA concentration of 0.5 μM.

At this experimental Mg^2+^ concentrations both *Model 1* and *Model 2* overlap. However, at lower Mg^2+^ concentrations, as shown in Figure 3, panels **E** and **F**, the two models give significantly different results. The presence of an alternative binding pathway in *Model 1* makes the folding in the presence of SAM less sensitive to the level of Mg^2+^, as opposed to *Model 2* in which the pseudoknot can only be formed in the presence of Mg^2+^. Several studies, however, have reported that at high concentrations of SAM most of the SAM-II RNA still form a pseudoknot and becomes folded, even in the absence of Mg^2+^ (6, 7). The latter findings are therefore consistent with *Model 1* and not with *Model 2*. Since *Model 1* also incorporates the previously discussed concentration dependent *Mg^2+^ Model* and its deduced parameters (10) (unlike *Models 3,4*,*6*), it is the only model that is qualitatively and quantitatively consistent with structural (population) changes observed in the absence of any ligands, after adding Mg^2+^ only, after adding SAM only and after adding both Mg^2+^ and SAM. Such data was collected using multiple techniques (4, 6–8).

Thus, *Model 1* provides the best explanation to the experimental qualitative and quantitative measurements for the SAM-II both at high (or in the presence) and at low (or in the absence) of Mg^2+^ concentrations, compared to the other models. Strikingly, the comparison between the models and experimental data illustrated that natural riboswitches exhibits the most efficient mechanism, at various levels of Mg^2+^, allowing the RNA to act as a highly efficient sensor and with a broad sensitivity-range.

Finally, the model is also consistent with the thermodynamically-driven tertiary structural changes in the presence of magnesium that were reported for other riboswitches (16, 18, 20–24). It is also consistent with the observed folding behavior for the thermodynamically driven translational riboswitches including the occurrence of tertiary interactions and the compaction of the riboswitch in the presence of both the ions and cognate ligand (25). Thus, in the following sections of this manuscript, only *Model 1* will be discussed further. A direct benefit of identifying the mechanism is that the titration curves for all RNA species involved in the model can be computed and studied in detail (Figure 4, panels **A** and **B)**. For the purpose of comparisons, the reader may refer to **Figure S2** for the corresponding plots for other models.

**Figure 4.**
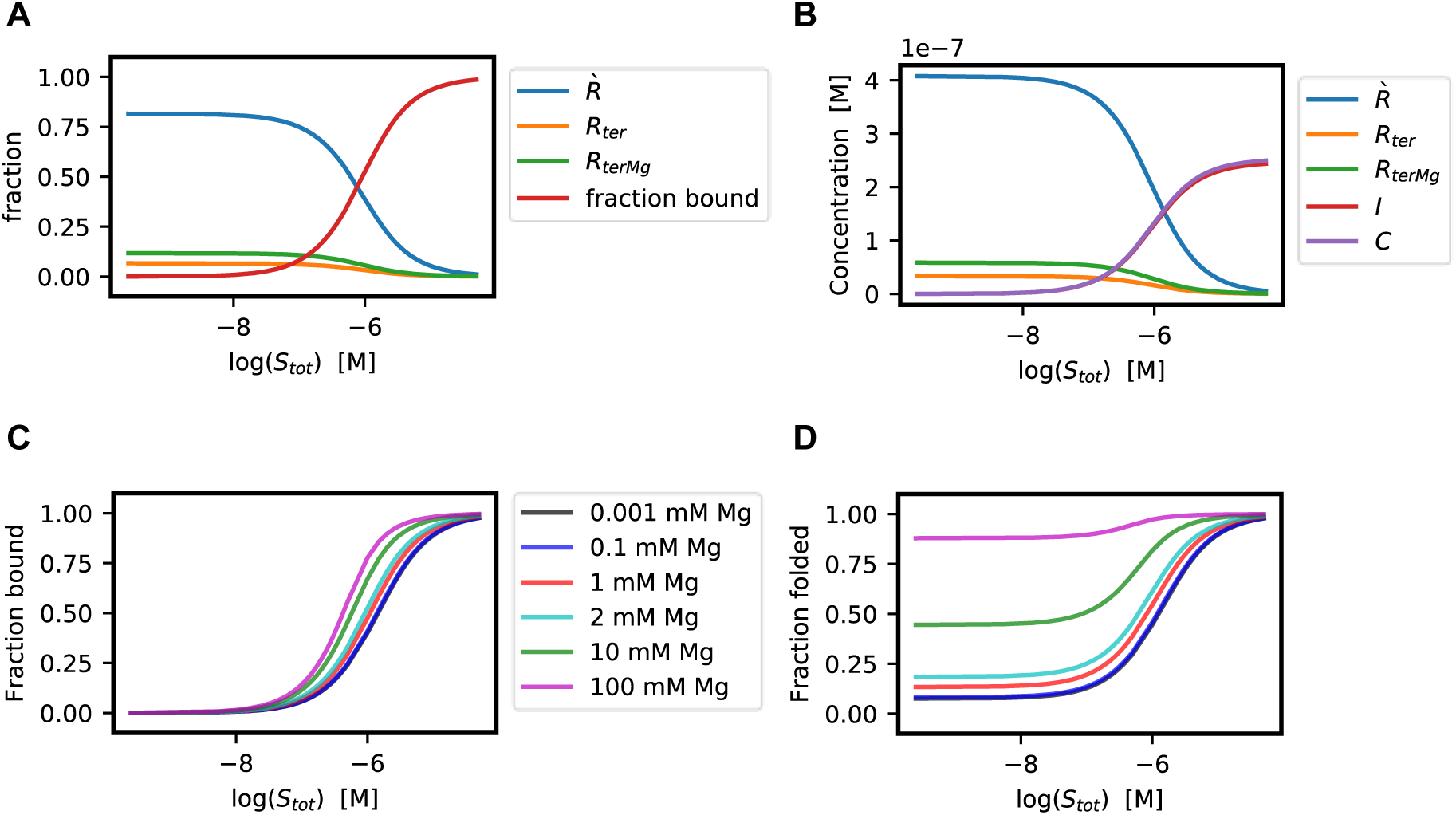
Titration curves for individual RNA species as a function of SAM concentration from *Model 1*. The plots in panels **A** and **B** illustrate the fractions and concentrations of the different RNA species computed from *Model 1*, as a function of total SAM concentration at a fixed total Mg^2+^ concentration of 2.0 mM and 0.5 μM RNA. Similar titration curves for individual RNA species computed based on this and other models discussed in this study (*Model 2* to *6*) are illustrated in **Figure S2**. The plots in panels **C** and **D** illustrate the *fraction bound* and *fraction folded* of the riboswitch as a function of SAM concentration, for different concentrations of Mg^2+^ ions, using the same model (*Model 1*). The latter titration curves illustrate that the magnesium level determines the sensitivity range of the riboswitch to its cognate ligand (SAM), by setting a threshold for the fraction of the already folded riboswitch.

### 3.3 The binding and folding mechanism of the SAM-II riboswitch

*Model 1*, which represents the riboswitch functional mechanism, shows that the aptamer pseudoknot of the SAM-II riboswitch, is initially stabilized by the magnesium ion to form a partially stable structure in an open conformation, paving the way for SAM to bind (Figure 1A). The binding of SAM then closes the binding pocket (forms a closed aptamer conformation, as observed in the crystal structure (4)) and further stabilizes the aptamer, essentially sequestering the ribosome binding site, and making it inaccessible for the Ribosome by a large free energy barrier. An alternative pathway for the riboswitch binding and folding may occur, in which SAM binds before magnesium. In either case, it is evident that the binding of both molecules is cooperative in nature, as the binding of one affects the affinity of the other (**Table 2**).

Since magnesium ions are usually present in the cell during transcription, the dominant product is dependent on the Mg^2+^ concentration (Figure 4C and D). In the presence of 2 mM magnesium, which was used in several *in vitro* experiments and also a relevant cell concentration (see (10) and references therein for a detailed discussion), approximately half of the SAM bound and folded population is also bound to Mg^2+^ (species *C*), while the other half is Mg^2+^ free (species *I*). This is illustrated in Figure 4B in which the latter two species are depicted in red and magenta, respectively. Thus, the fact that the crystal structure of the SAM- II riboswitch did not show a bound Mg^2+^ ion (4), whereas a very similar SAM-V riboswitch (26) showed a bound Mg^2+^ ion, can be explained by the notion that both states co-exist, even in the abundance of SAM.

As shown previously based on Mg^2+^ data, the formation of the *R*_*ter*_ is not favorable, with a free energy of 1.483 kcal/mol. *R*_*ter*_, however, can be stabilized by the binding of Mg^2+^ to form *R*_*terMg*_ with a binding energy of -4.016 kcal/mol. SAM can bind to both *R*_*ter*_ and *R*_*terMg*_ with a stabilizing binding energies of -9.608 and -9.289 kcal/mol, respectively, to form *I* and *C*, which represent the open and closed folded aptamer conformations. Finally, a Mg^2+^ ion may further stabilize *I* (the open conformation) with a free energy of -3.696 kcal/mol by zipping the binding pocket, converting it to the closed conformation *C*. The expected SAM binding using single molecule FS (*dG*_*FS*_) based on *Model 1* is −9.425 kcal/mol, which is very close to the experimentally reported value −9.350 kcal/mol at room temperature. Notably, this model (*Model 1*) seems to be the most efficient compared to the other models (Figure 3A). That is, the fraction folded (i.e. forming a pseudoknot and hence essentially sequestering the ribosome binding site) calculated using *Model 1* at relevant (near physiological) concentrations of SAM and Mg^2+^, reaches a level that is higher than other models. The latter suffer from the fact that a significant fraction of the RNA is not responsive to the SAM (i.e. not functional) at these substrate levels.

### 3.4 Magnesium concentration determines the sensitivity range of the riboswitch to the its congnate ligand

Interestingly, the concentration of Mg^2+^ achieves a certain level of folding prior to SAM binding, allowing the pre-nucleated pseudoknot-containing riboswitch to strongly respond to SAM only if the latter exceeds a certain threshold concentration. Such property is probably to be desirable in the cell and had been often discussed in transcriptional or kinetically driven riboswitches (27, 28). In other words, Mg^2+^ controls the sensitivity range of the riboswitch to the primary ligand. Figures 4C and **4D** show the *fraction folded* and *fraction bound* of the RNA (simulated using *Model 1*) as a function of SAM for different Mg^2+^ levels. It is evident that the sensitivity range of the riboswitch decreases as Mg^2+^ ions increase.

## 4 DISCUSSION

### 4.1 Prediction of the fraction of riboswitch folded as a function of both Mg^2+^ and SAM: constructing a free energy landscape

The determined binding and folding free energies (which are the model parameters) allow us to construct a standard binding and folding free energy landscape (Figure 5), which provides insights into the system, at standard conditions. Such energies can be thought of as the molar free energy differences, at standard room temperature and pressure, and at saturating concentrations of the ligands. It is evident from the figure that one can determine the standard free energy difference between any two states of the system, for example *R* and *I*, even if the conversion between these states must go through one or more intermediate.

**Figure 5.**
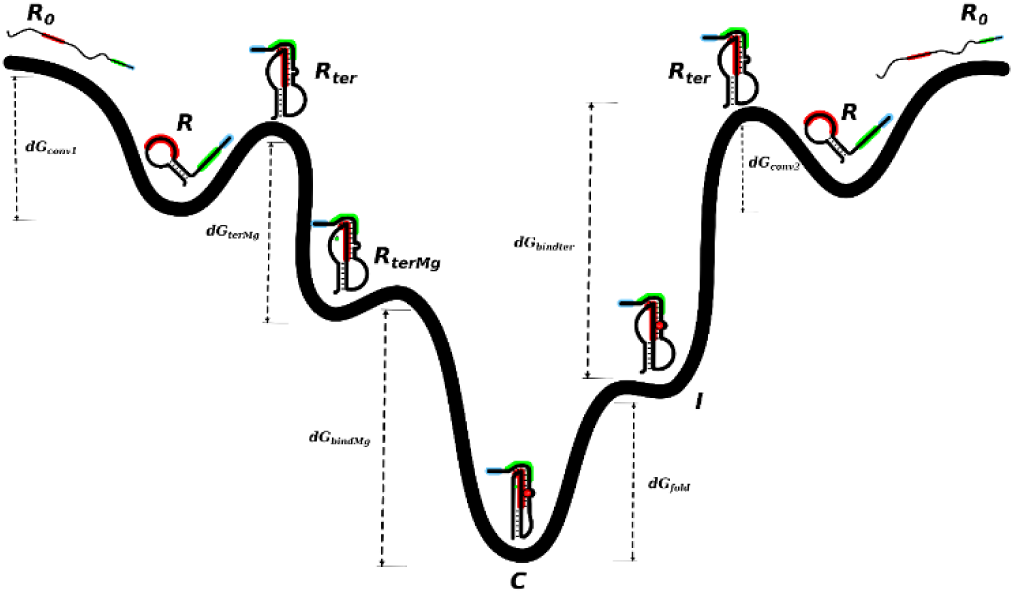
The free energy landscape of the SAM-II riboswitch. The figure illustrates the free energy landscape of the SAM-II riboswitch with corresponding standard free energy differences between a give state and the neighbouring one are shown on the arrows. The initial unfolded or incorrectly folded state (*R*_0_) here acts as the reference. The values of the free energy differences shown on the figure are tabulated in Table 2 and are further discussed in the text.

More importantly, the model can be used to quantitatively predict the probabilities of all species in the ensemble at varying concentrations of two or three reactants. For example, the three-dimensional (3D) titration curves in **Figures S3** and **S4,** respectively, illustrate the changes of the various fractions (or concentrations) of RNA species as a function of 1) SAM and Mg^2+^ concentrations, at a fixed total RNA concentration, and 2) total RNA and Mg^2+^ concentrations, at a fixed total SAM concentration. This study therefore shows a case where concentration dependent mechanistic/mathematical models were combined with conformer populations (ensembles calculations based on secondary structure energetics) and linked to multiple experimental ligand binding studies to provide a rich and detailed understanding of the system. It is worth noting that the methods described can also be used to compute ligand titration curves at different transcription lengths. More generally, when basic individual conformer probabilities along with models for single or multiple Mg^2+^ ions (10, 19) are considered, it is possible to directly apply or adapt the mathematical framework of the models in this study to any RNA system (including other riboswitches) that undergoes structural changes upon ligand binding.

### 4.2 Limitations

The presented theoretical framework described come with several limitations. Most importantly, the models were not aimed to describe kinetic effects, which are often seen for cognate ligand effect on transcriptional riboswitches. Nonetheless, this work can be used as basis for developing such effects. Moreover, many riboswitches involve the binding to multiple ions and can be more thoroughly described by the folding of multiple tertiary elements. Hence, future work needs to illustrate the implementation of the framework used here on more complex systems.

## 5 CONCLUSIONS

This study deconvolutes the mechanism by which the SAM-II riboswitch functions. In this mechanism, RNA with a pre-existing pseudoknot binds tightly with SAM, in the presence or the absence of magnesium ions, and hence become more stable, eventually dominating the RNA population. The level of magnesium ions sets the concentration threshold and range in which the riboswitch can be responsive to SAM. It is evident that, such mechanism may be similar for other riboswitches, paving the way for practical and novel avenues in drug design. The presented analytical/mathematical framework, based on linked equilibria model, which combines secondary structure energetics with ligand binding, though specifically used here to quantitatively model the SAM-II riboswitch, is yet applicable to other riboswitches. Moreover, the presented models and methods that failed to describe the behavior of this riboswitch, may still be generally useful and applicable to other RNA that undergoes conformational changes in the presence of one or more ligands. We anticipate that the mechanisms and tools, contributed via this work will advance our understanding of RNA function.

## 6 SUPPORTING INFORMATION

Supporting Information including Supplementary methods, figures and tables are available online.

## 7 CONFLICT OF INTERESTS

The author declares that he has shares in Biocomplexity for Research and Consulting, which is a for-profit company.

## Notes

### Summary of Updates

The manuscript was re-arranged, and parts were re-written to become more concise. Detailed derivations were moved to the SI. Main equations were summarized in a table for clarity. Affiliation was updated to more clearly reflect current address and where the work was performed.

