## Supplementary Material for "Modeling Translational Riboswitches: The impact of SAM concentration on the folding of the SAM-II riboswitch"

Osama Alaidi [1,2]

[1] Biocomplexity for Research and Consulting, Cairo, Egypt.

[2] **Current address:** Department of Pharmaceutical Sciences, University of Tennessee Health Science Center, Memphis, TN, 38163, United States.

ORCID: Osama Alaidi: 0000-0002-1292-6789

### Contents


1 Supplementary methods ... 3
  1.1 *Model 0*: Predicting fractions of RNA conformers in the absence of ligands ... 3
  1.2 *Model 1* ... 5
  1.3 *Model 2* ... 12
  1.4 *Model 3* ... 15
  1.5 *Model 4* ... 22
  1.6 *Model 5* ... 23
  1.7 *Model 6* ... 30
2 Supplementary Figures ... 36
3 Supplementary Tables ... 40
4 References ... 43

### 1 Supplementary methods

The following section is aimed to illustrate the derivation of expressions for the concentration of the final state, which is the bound and folded RNA (designated by $I$ or $C$, depending on the model). A concise summary of these expressions (for *Models 1-6*) is shown in **Table S2**. For each model, the derivation starts by defining the equilibrium constants for the model, and ends with a solution for a quadratic equation in which its coefficients are functions of the equilibrium constants and the total concentrations of the reacting species ($R_{tot}$ , $S_{tot}$ and $Mg_{tot}$). Pictorial representations of these models are illustrated in **Figures 1A-1F** in the main text. Further, all species and constants are defined in the **Methods** section of the main text. In all models here, the free $Mg^{2+}$ions concentration here is assumed to be equal to the total $Mg^{2+}$ concentrations (i.e. $Mg \approx Mg_{tot}$), see the detailed discussion about this approximation in reference (1).

#### 1.1 *Model 0*: Predicting fractions of RNA conformers in the absence of ligands

**Figure S1** shows a model for RNA conversion and folding in the absence of both magnesium and the cognate ligand (SAM). In the latter, the following quantities are defined as follows.

The initial conditions are represented by $R_{tot}$ where, $R_{tot}$ is the total RNA concentration. The equilibrium constants are defined as, $K_{conv1}$ and $K_{conv2}$ which are the equilibrium constants for conversion and pseudoknot folding, respectively. $K_{eq}$ is the overall equilibrium for *Model 0*. $R_0$ is the concentration of the RNA in the non-reactive state (not forming a plausible secondary structure). $R$ is the concentration of the RNA with a secondary structure that can proceed to form the pseudoknot. At equilibrium we have,

$K_{conv1} = \frac{R}{R_0}$ , hence $R = K_{conv1} R_0$

and $K_{conv2} = \frac{R_{ter}}{R}$, hence,

$$\begin{aligned} R_{ter} &= K_{conv2}\, R \\ &= K_{conv2}\, K_{conv1} R_0 \\ &= K_{eq} R_0 \end{aligned} \qquad \text{Eq. 0.1}$$

$K_{conv1}$ and $K_{conv2}$ can be obtained from ratios between their probabilities via secondary structure calculations using the substructure search method described in (2), *where*,

$$K_{conv2} = \frac{P_{R_{ter}}}{P_R} \qquad \text{Eq. 0.2}$$

and

$$K_{conv1} = \frac{P_R}{P_{R0}} \qquad \text{Eq. 0.3}$$

Using conservation of moieties and a series of substitutions we get:

$$\begin{aligned} R_{ter} &= K_{eq} R_0 \\ &= \frac{K_{eq}}{1+K_{eq}} (1 - P_R) \times R_{tot} \\ &= R_{tot} \Big/ \left(1 + \frac{1}{K_{eq}} + \frac{1}{K_{conv2}}\right) \end{aligned} \qquad \text{Eq. 0.4}$$

Equivalently,

$$R_{ter} = K_{ter}\, R^{`} \qquad \text{Eq. 0.5}$$

### 1.2 *Model 1*

According to the mass balance law and from the definition of equilibrium and **Figure 1A**, the following can be defined,

$$K_{ter} = \frac{R_{ter}}{R^`} \qquad \text{Eq. 1.1}$$

$$K_{Mg} = \frac{R_{terMg}}{R_{ter} \times Mg} \qquad \text{Eq. 1.2}$$

$$K_{bindter} = \frac{I}{R_{ter} \times S} \qquad \text{Eq. 1.3}$$

$$K_{bindMg} = \frac{C}{R_{terMg} \times S} \qquad \text{Eq. 1.4}$$

$$K_{fold} = \frac{C}{I \times Mg} \qquad \text{Eq. 1.5}$$

From the definition of equilibrium above (equations 1.1 - 1.5), we have,

$$C = K_{bindMg} \times R_{terMg} \times S$$

$$= K_{bindMg} \times K_{Mg} \times R_{ter} \times S \times Mg$$

By substituting for $R_{ter}$ and $R_{terMg}$, from Eq. 1.1 and Eq. 1.2, we get

$$= R^` \times S \times Mg \times (K_{ter} \times K_{bindMg} \times K_{Mg})$$

$$C = R^` \times S \times \boldsymbol{\alpha} \qquad \text{Eq. 1.6}$$

*where*, $\boldsymbol{\alpha} = (K_{ter} \times K_{bindMg} \times K_{Mg}) \times Mg$

From the conservation of moieties, the total RNA is equal to,

$$R_{tot} = R^` + R_{ter} + R_{terMg} + I + C \qquad \text{Eq. 1.7}$$

By substitution of equations 1.1, 1.2, 1.5, in equation 1.7,

$$R_{tot} = R^` + R^` K_{ter} + R_{ter}K_{Mg}.Mg + \frac{C}{K_{fold}.Mg} + C$$

$$= R^{`} + R^{`}K_{ter} + R^{`}K_{ter}K_{Mg}.Mg + C\left(\frac{1}{K_{fold}.Mg} + 1\right)$$

$$R_{tot} = R^{`} + R^{`}K_{ter} + R^{`}K_{ter}K_{Mg}.Mg + C\beta \qquad \text{Eq. 1.8}$$

*where,*

$$\beta = \left(\frac{1}{K_{fold}.Mg} + 1\right) \qquad \text{Eq. 1.9}$$

Thus,

$$R^{`} + R^{`}K_{ter} + R^{`}K_{ter}K_{Mg}.Mg = R_{tot} - C\beta$$

$$R^{`} = \frac{R_{tot} - C\beta}{1 + K_{ter} + K_{ter}.K_{Mg}.Mg} = \frac{R_{tot} - C\beta}{\gamma} \qquad \text{Eq. 1.10}$$

And

$$\gamma = 1 + K_{ter} + K_{ter}.K_{Mg}.Mg = 1 + K_{ter}(1 + K_{Mg}.Mg) \qquad \text{Eq. 1.11}$$

The total SAM concentration $S_{tot}$ , is a conserved quantity and is equal to

$$S_{tot} = S + I + C \qquad \text{Eq. 1.12}$$

Thus, the free SAM is described by,

$$S = S_{tot} - (I + C)$$

$$= S_{tot} - \left(\frac{C}{K_{fold}.Mg} + C\right)$$

$$= S_{tot} - C\left(\frac{1}{K_{fold}.Mg} + 1\right)$$

$$S = S_{tot} - C\beta \qquad \text{Eq. 1.13}$$

To obtain the concentrations of the bound aptamer, $C$, in terms of the equilibrium constants and the total concentration of the reactants, $R_{tot}$ and $S_{tot}$ , we substitute for $R^{`}$ and $S$ from equations 1.10 and 1.13 in Eq. 1.6,

$C = R^{`} \times S \times \boldsymbol{\alpha}$

$$= \left(\frac{R_{tot} - C\beta}{\gamma}\right) \times (S_{tot} - C\beta) \times \boldsymbol{\alpha}$$

$$= (R_{tot} - C\beta) \times (S_{tot} - C\beta) \times \frac{\boldsymbol{\alpha}}{\gamma}$$

Thus,

$$\frac{\gamma}{\boldsymbol{\alpha}} C = (R_{tot} - C\beta) \times (S_{tot} - C\beta)$$

By distributing the brackets, subtracting $\frac{\gamma}{\boldsymbol{\alpha}} C$ from both sides and taking $C$ as a common factor, we get a quadratic,

$$\beta^2 C^2 - R_{tot}\beta C - S_{tot}\beta C - \frac{\gamma}{\boldsymbol{\alpha}} C + S_{tot}R_{tot} = 0$$

$$(\beta^2)C^2 + \left(-\beta(R_{tot} + S_{tot}) - \frac{\gamma}{\boldsymbol{\alpha}}\right) C + (S_{tot}R_{tot}) = 0$$

The solution to the quadratic for $C$ is,

$$C = \frac{-b \pm \sqrt{b^2 - 4ac}}{2a} \qquad \text{Eq. 1.14}$$

*Where,*

$$a = \beta^2$$

$$b = -\beta(R_{tot} + S_{tot}) - \frac{\gamma}{\boldsymbol{\alpha}}$$

$$c = S_{tot}R_{tot}$$

Hence, a final expression for computing $C$ is given by (Eq. 1.15),

$$C = \frac{-\left(-\beta(R_{tot} + S_{tot}) - \frac{\gamma}{\boldsymbol{\alpha}}\right) \pm \sqrt{\left(-\beta(R_{tot} + S_{tot}) - \frac{\gamma}{\boldsymbol{\alpha}}\right)^2 - 4\beta^2(S_{tot}R_{tot})}}{2\beta^2}$$

The above expression describes the concentration of the bound riboswitch with a pseudoknot as a function of the total concentrations of reactants, $R_{tot}$, $Mg$ and $S_{tot}$ and the equilibrium constants.

***Determination of Model 1 parameters***

The overall (apparent) dissociation constant which was determined experimentally using Isothermal calorimetry (ITC) and reported by Gilbert *et al* (3). describes the following reaction,

$$Unbound\ RNA + SAM \underset{Ka_{ITC}}{\Longleftrightarrow} bound\ RNA$$

$$\left[R_0 + R + R_{ter} + R_{terMg}\right] + [S] \underset{Ka_{ITC}}{\Longleftrightarrow} [I + C]$$

$$\left[R^{`} + R_{ter} + R_{terMg}\right] + [S] \underset{Ka_{ITC}}{\Longleftrightarrow} [I + C]$$

$Kd_{ITC}$ was experimentally determined to be $0.67 \pm 0.06\ \mu M$. Hence, this experimental dissociation constant $Kd_{ITC} = \frac{1}{Ka_{ITC}}$, describes the following equilibrium,

$$Ka_{ITC} = \frac{Bound\ RNA}{Unbound\ RNA\ \times free\ SAM}$$

$$= \frac{[I + C]}{\left[R_0 + R + R_{ter} + R_{terMg}\right] \times [S]}$$

$$= \frac{[I + C]}{\left[R^{`} + R_{ter} + R_{terMg}\right] \times [S]}$$

$$= \frac{\left[\left(\frac{1}{K_{fold}.Mg}\right).C + C\right]}{\left[\frac{R_{ter}}{K_{ter}} + \frac{R_{terMg}}{K_{Mg}.Mg} + R_{terMg}\right] \times [S]}$$

$$= \frac{C\left[\left(\frac{1}{K_{fold}.Mg}\right) + 1\right]}{R_{terMg} \times S \times \left[\frac{1}{K_{ter}K_{Mg}.Mg} + \frac{1}{K_{Mg}.Mg} + 1\right]}$$

$$= \frac{C \times \beta_1}{R_{terMg} \times S \times \alpha_1}$$

Thus,

$$Ka_{ITC} = K_{bindMg}\ \frac{\beta_1}{\alpha_1} \qquad \text{Eq. 1.16}$$

*where,* $K_{bindMg}$ is the SAM binding association constant of $R_{terMg}$, $\beta_1 = \left(\frac{1}{K_{fold}.Mg}\right) + 1$ and, $\alpha_1 = \left(\frac{1}{K_{ter}K_{Mg}.Mg} + \frac{1}{K_{Mg}.Mg} + 1\right)$. The value of $\alpha_1$ is given and depends on the $Mg^{2+}$ ions concentration, whereas $\beta_1$ is unknown and cannot be determined from the thermodynamic cycle alone.

Similarly (and alternatively),

$$Ka_{ITC} = \frac{Bound\ RNA}{Unbound\ RNA\ \times free\ SAM}$$

$$= \frac{[I + C]}{[R_0 + R + R_{ter} + R_{terMg}] \times [S]}$$

$$= \frac{[I + C]}{[R^{`} + R_{ter} + R_{terMg}] \times [S]}$$

$$= \frac{[I + K_{fold}.Mg.I]}{\left[\frac{R_{ter}}{K_{ter}} + R_{ter} + K_{Mg}.Mg.R_{ter}\right] \times [S]}$$

$$= \frac{I\left[K_{fold}.Mg + 1\right]}{R_{ter} \times S \times \left[\frac{1}{K_{ter}} + K_{Mg}.Mg + 1\right]}$$

$$= \frac{I \times \beta_2}{R_{ter} \times S \times \alpha_2}$$

Thus,

$$Ka_{ITC} = K_{bindter}\ \frac{\beta_2}{\alpha_2} \qquad \text{Eq. 1.17}$$

*where,* $K_{bindter}$ is the SAM binding association constant of $R_{ter}$, $\alpha_2 = \left(\frac{1}{K_{ter}} + K_{Mg}\ .Mg + 1\right)$ and $\beta_2 = (K_{fold}\ .Mg + 1)$.

The previously reported FS studies monitor RNA folding in the presence of the ligand relative to its absence (4). In other words, the apparent binding constant determines the fraction of RNA species with a pseudoknot and bound to the ligand compared to the fraction with a pseudoknot but without the ligand. Therefore, binding constants from such FS studies, represent the following reaction,

$$Folded\ \&\ unbound\ RNA + SAM \underset{Ka_{FS}}{\Longleftrightarrow} Folded\ \&\ bound\ RNA$$

$$[R_{ter} + R_{terMg}] + [S] \underset{Ka_{FS}}{\Longleftrightarrow} [I + C]$$

$$Ka_{FS} = \frac{Bound\ RNA\ species\ with\ a\ pseudoknot}{Unbound\ RNA\ species\ with\ a\ pseudoknot\ \times free\ SAM}$$

$$= \frac{[I + C]}{[R_{ter} + R_{terMg}] \times [S]}$$

$$= \frac{[I + K_{fold}.Mg.I]}{[R_{ter} + K_{Mg}.Mg.R_{ter}] \times [S]}$$

$$= \frac{I\,[K_{fold}.Mg + 1]}{R_{ter} \times S \times [K_{Mg}.Mg + 1]}$$

$$= \frac{I \times \beta_2}{R_{ter} \times S \times \epsilon}$$

*Thus,*

$$Ka_{FS} = K_{bindter}\ \frac{\beta_2}{\epsilon} \qquad \text{Eq. 1.18}$$

*where,* $K_{bindter}$ is the SAM binding association constant of $R_{ter}$, $\epsilon = (K_{Mg}\ .Mg + 1)$ and $\beta_2 = (K_{fold}\ .Mg + 1)$.

For convenience and validation, by substitution for $\beta_2$ (from Eq. 1.17) into Eq. 1.18, a relation between $Ka_{FS}$ and $Ka_{ITC}$ can be established,

$$Ka_{FS} = K_{bindter}\ \frac{\beta_2}{\epsilon} = K_{bindter} \frac{Ka_{ITC}\ \alpha_2}{\epsilon\ K_{bindter}} = Ka_{ITC}\ \frac{\alpha_2}{\epsilon} \qquad \text{Eq.1.19}$$

Since, $K_{Mg}$and $K_{ter}$ is known from magnesium data (1), $\alpha_2$ and $\epsilon$ can be computed, and hence knowing one of the two constants, either $Ka_{ITC}$ or $Ka_{FS}$ can determine the other. Each of the latter two constants are determined by two of the unknown constants.

To determine each of $K_{fold}$, $K_{bindter}$ and $K_{bindMg}$ a third relation from the thermodynamic cycle is needed. From the cycle in **Figure 2A**, we have the following,

$$\Delta G_{fold} + \Delta G_{bindter} - \Delta G_{bindMg} - \Delta G_{Mg} = 0$$

$$\Delta G_{fold} = \Delta G_{bindMg} + \Delta G_{Mg} - \Delta G_{bindter}$$

By dividing both sides by $-RT$ and taking the exponents of both sides, we obtain,

$$e^{\frac{-\Delta G_{fold}}{RT}} = e^{\frac{-\Delta G_{bindMg}}{RT} + \frac{-\Delta G_{Mg}}{RT} - \frac{-\Delta G_{bindter}}{RT}}$$

$$e^{\frac{-\Delta G_{fold}}{RT}} = \frac{e^{\frac{-\Delta G_{bindMg}}{RT}} \;.\; e^{\frac{-\Delta G_{Mg}}{RT}}}{e^{\frac{-\Delta G_{bindter}}{RT}}}$$

$$K_{fold} = \frac{K_{bindMg}}{K_{bindter}} K_{Mg}$$

By substitution of $K_{bindMg}$ (from Eq. 1.16) and $K_{bindter}$ (from Eq. 1.18) in terms of experimentally measured constants ($Ka_{ITC}$ and $Ka_{FS}$), we obtain,

$$K_{fold} = \frac{Ka_{ITC}\ \alpha_1\ \beta_2}{Ka_{FS}\ \epsilon\ \beta_1} K_{Mg}$$

$$K_{fold}\frac{\beta_1}{\beta_2} = \frac{Ka_{ITC}\ \alpha_1\ K_{Mg}}{Ka_{FS}\ \epsilon} = \frac{\sigma}{\rho}$$

Then by the substitution for $\beta_2$ and $\beta_1$ (as defined in Eq. 1.16 and Eq. 1.17), and solving for $K_{fold}$, the latter constant can be determined,

$$K_{fold} = \frac{\frac{\sigma}{Mg} - \rho}{\rho Mg - \sigma} \qquad \text{Eq. 1.20}$$

Where, $\sigma = Ka_{FS}\ \epsilon$ and $\rho = Ka_{ITC}\ \alpha_1\ K_{Mg}$ . Once $K_{fold}$ is determined from Eq. 1.20, $K_{bindMg}$ and $K_{bindter}$ can be determined through Eq. 1.16 - 1.18.

### 1.3 *Model 2*

This model (illustrated in **Figure 1B**) is similar to the *Model 1* but differs in that $Mg^{2+}$ is essential for SAM binding. The final conformer is denoted as $C$. A summary and final equations are reviewed below.

From the mass conservation law and **Figure 1B**, the following can be defined,

$$K_{ter} = \frac{R_{ter}}{R^{`}} \quad \text{Eq. 2.1}$$

$$K_{Mg} = \frac{R_{terMg}}{R_{ter} \times Mg} \quad \text{Eq. 2.2}$$

$$K_{bindMg} = \frac{C}{R_{terMg} \times S} \quad \text{Eq. 2.3}$$

$$C = K_{bindMg} \times R_{terMg} \times S = K_{bindMg} .K_{ter} .K_{Mg} .Mg.R^{`}.S = R^{`}.S.\boldsymbol{\alpha} \quad \text{Eq. 2.4}$$

Where, $\boldsymbol{\alpha} = K_{bindMg} .K_{ter} .K_{Mg} .Mg$. Taking into account that $S = S_{tot} - C$ and $R_{tot} = R^{`} + R_{ter} + R_{terMg} + C$ (by substituting the equilibrium constants), will give expressions similar to Eq. 1.13 and Eq. 1.10 with $\beta = 1$, that is,

$$R^{`} = \frac{R_{tot} - C}{1 + K_{ter} + K_{ter} .K_{Mg}.Mg} = \frac{R_{tot} - C}{\gamma} \quad \text{Eq. 2.5}$$

Where, $\gamma = 1 + K_{ter} + K_{ter} .K_{Mg}.Mg$

Substituting for $S$ and $R^{`}$ in equation 2.4 and solving for the species $C$ will lead to a quadratic similar to the one obtained for *Model 1* (Eq. 1.15) but with $\beta = 1$, which is written as,

$$C = \frac{-\left(-(R_{tot} + S_{tot}) - \frac{\gamma}{\boldsymbol{\alpha}}\right) \pm \sqrt{\left(-(R_{tot} + S_{tot}) - \frac{\gamma}{\boldsymbol{\alpha}}\right)^2 - 4(S_{tot}R_{tot})}}{2} \quad \text{Eq. 2.6}$$

The quadratic equation has two possible solutions for $C$. Only one of the two roots is physically sensible, and can represent concentrations (i.e. positive real number, with a maximum value equal or less than $R_{tot}$).

***Determination of Model 2 parameters***

Model parameters can be obtained in a similar way to the previous model except that the intermediate $I$ is absent i.e. $I = 0$ and hence the factor $\beta_1 = 1$. Briefly, the overall (apparent) dissociation constant which was determined experimentally using Isothermal calorimetry (ITC), and reported by Gilbert *et al.* (3) describes the following reaction,

$$Unbound\ RNA + SAM \underset{Ka_{ITC}}{\Longleftrightarrow} bound\ RNA$$

$$\left[R_0 + R + R_{ter} + R_{terMg}\right] + [S] \underset{Ka_{ITC}}{\Longleftrightarrow} [C]$$

$$\left[R^{`} + R_{ter} + R_{terMg}\right] + [S] \underset{Ka_{ITC}}{\Longleftrightarrow} [C]$$

$Kd_{ITC}$ was experimentally determined to be $0.67 \pm 0.06\,\mu M$. Since this experimental dissociation constant $Kd_{ITC} = \frac{1}{Ka_{ITC}}$, describes the following equilibrium,

$$Ka_{ITC} = \frac{Bound\ RNA}{Unbound\ RNA\ \times free\ SAM}$$

$$= \frac{[C]}{\left[R_0 + R + R_{ter} + R_{terMg}\right] \times [S]}$$

$$= \frac{[C]}{\left[R^{`} + R_{ter} + R_{terMg}\right] \times [S]}$$

$$= \frac{[C]}{\left[\frac{R_{ter}}{K_{ter}} + \frac{R_{terMg}}{K_{Mg}.Mg} + R_{terMg}\right] \times [S]}$$

$$= \frac{C}{R_{terMg} \times S \times \left[\frac{1}{K_{ter}K_{Mg}.Mg} + \frac{1}{K_{Mg}.Mg} + 1\right]}$$

$$= \frac{C}{R_{terMg} \times S \times \alpha_1}$$

Thus,

$$Ka_{ITC} = \frac{K_{bindMg}}{\alpha_1} \qquad \text{Eq. 2.7}$$

*where,* $K_{bindMg}$ is the SAM binding association constant of $R_{terMg}$, and $\alpha_1 = \left(\frac{1}{K_{ter}K_{Mg}.Mg} + \frac{1}{K_{Mg}.Mg} + 1\right)$. The value of $\alpha_1$ is given and it depends on the $Mg^{2+}$ ions concentration along with other equilibrium constants.

As in the previous model, again the reported FS studies monitor RNA folding in the presence of the ligand relative to its absence. That is, the apparent binding constant determines the fraction of RNA species with a pseudoknot and bound to the ligand compared to the fraction with a pseudoknot but without the ligand considering the ligand concentration. Therefore, if this model is true, the binding constants from FS studies, would represent the following reaction,

$$Folded\ \&\ unbound\ RNA + SAM \xLeftrightarrow[Ka_{FS}]{} Folded\ \&\ bound\ RNA$$

$$[R_{ter} + R_{terMg}] + [S] \xLeftrightarrow[Ka_{FS}]{} [I + C]$$

$$Ka_{FS} = \frac{Bound\ RNA\ species\ with\ a\ pseudoknot}{Unbound\ RNA\ species\ with\ a\ pseudoknot\ \times free\ SAM}$$

$$= \frac{[C]}{[R_{ter} + R_{terMg}] \times [S]}$$

$$= \frac{[C]}{\left[\frac{R_{terMg}}{K_{Mg}.Mg} + R_{terMg}\right] \times [S]}$$

$$= \frac{[C]}{R_{terMg} \times S \times \left[\frac{1}{K_{Mg}.Mg} + 1\right]}$$

$$Ka_{FS} = K_{bindMg} \frac{1}{\left[\frac{1}{K_{Mg}.Mg} + 1\right]} = K_{bindMg} \cdot \gamma \qquad \text{Eq. 2.8}$$

Thus, for the purpose of validation, a relation between $Ka_{FS}$ and $Ka_{ITC}$ can be established,

$$Ka_{FS} = Ka_{ITC} \cdot \alpha_1 \cdot \gamma \qquad \text{Eq. 2.9}$$

*where,* $\gamma = \frac{1}{\left[\frac{1}{K_{Mg}.Mg} + 1\right]}$.

### 1.4 *Model 3*

As in the previous model, from the definition of equilibrium and **Figure 1C**, the following can be defined,

$$K_{conv1} \quad = \frac{R}{R_0} \qquad \text{Eq. 3.1}$$

$$K_{terE} \quad = \frac{R_{terE}}{R \times Mg} \qquad \text{Eq. 3.2}$$

$$K_{bind} \quad = \frac{I}{R \times S} \qquad \text{Eq. 3.3}$$

$$K_{bindE} = \frac{C}{R_{terE} \times S} \qquad \text{Eq. 3.4}$$

$$K_{fold} \quad = \frac{C}{I \times Mg} \qquad \text{Eq. 3.5}$$

Note that, $K_{bindE} = \frac{C}{R_{terE} \times S} = \frac{K_{fold}\, I\, Mg}{K_{terE}\, R\, Mg} = \frac{K_{fold}.I\,.Mg}{K_{terE}\,.R\,.Mg\,.S} = \frac{K_{fold}.K_{bind}\,.\,R\,.\,S\,.Mg}{K_{terE}\,.R\,.Mg\,.S} = \frac{K_{fold}.K_{bind}}{K_{terE}}$

From the definition of equilibrium we have,

$$C = K_{bindE} \times R_{terE} \times S$$

$$= K_{bindE} \times K_{terE} \times R \times S \times Mg$$

By substituting for $R$, from Eq. 3.1, we get

$$= R_0 \times S \times Mg \times (K_{conv1} \times K_{bindE} \times K_{terE})$$

$$C = R_0 \times S \times \boldsymbol{\alpha} \qquad \text{Eq. 3.6}$$

*where*, $\boldsymbol{\alpha} = (K_{conv1} \times K_{bindE} \times K_{terE}) \times Mg$

From the conservation of moieties, the total RNA is equal to,

$$R_{tot} = R_0 + R + R_{terE} + I + C \qquad \text{Eq. 3.7}$$

By substitution of equations 3.1, 3.2, 3.5, in equation 3.7,

$$R_{tot} \;=\; R_0 + R_0 K_{conv1} + R K_{terE}.Mg + \frac{C}{K_{fold}.Mg} + C$$

$$= R_0 + R_0K_{conv1} + R_0K_{conv1}K_{terE}.Mg + C\left(\frac{1}{K_{fold}.Mg} + 1\right)$$

$$R_{tot} = R_0 + R_0K_{conv1} + R_0K_{conv1}K_{terE}.Mg + C\beta \qquad \text{Eq. 3.8}$$

*where,*

$$\beta = \left(\frac{1}{K_{fold}.Mg} + 1\right) \qquad \text{Eq. 3.9}$$

Thus,

$$R_0 + R_0K_{conv1} + R_0K_{conv1}K_{terE}.Mg = R_{tot} - C\beta$$

$$R_0 = \frac{R_{tot} - C\beta}{1 + K_{conv1} + K_{conv1}\,K_{terE}.Mg} = \frac{R_{tot} - C\beta}{\gamma} \qquad \text{Eq. 3.10}$$

And

$$\gamma = 1 + K_{conv1} + K_{conv1}\,K_{terE}.Mg = 1 + K_{conv1}(1 + K_{terE}.Mg) \qquad \text{Eq. 3.11}$$

The total SAM concentration $S_{tot}$ , is a conserved quantity and is equal to

$$S_{tot} = S + I + C \qquad \text{Eq. 3.12}$$

Thus, the free SAM is described by,

$$S = S_{tot} - (I + C)$$

$$= S_{tot} - \left(\frac{C}{K_{fold}.Mg} + C\right)$$

$$= S_{tot} - C\left(\frac{1}{K_{fold}.Mg} + 1\right)$$

$$S = S_{tot} - C\beta \qquad \text{Eq. 3.13}$$

To obtain the concentrations of the bound aptamer, $C$, in terms of the equilibrium constants and the total concentration of the reactants, $R_{tot}$ and $S_{tot}$ , we substitute for $R_0$ and $S$ from equations 3.10 and 3.13 in eq. 3.6,

$$C = R_0 \times S \times \boldsymbol{\alpha}$$

$$= \left(\frac{R_{tot} - C\beta}{\gamma}\right) \times (S_{tot} - C\beta) \times \boldsymbol{\alpha}$$

$$= (R_{tot} - C\beta) \times (S_{tot} - C\beta) \times \frac{\boldsymbol{\alpha}}{\gamma}$$

Thus, $\frac{\gamma}{\boldsymbol{\alpha}}C = (R_{tot} - C\beta) \times (S_{tot} - C\beta)$

By distributing the brackets, subtracting $\frac{\gamma}{\boldsymbol{\alpha}}C$ from both sides and taking $C$ as a common factor, we get a quadratic,

$$\beta^2C^2 - R_{tot}\beta C - S_{tot}\beta C - \frac{\gamma}{\boldsymbol{\alpha}}C + S_{tot}R_{tot} = 0$$

$$(\beta^2)C^2 + \left(-\beta(R_{tot} + S_{tot}) - \frac{\gamma}{\boldsymbol{\alpha}}\right)C + (S_{tot}R_{tot}) = 0$$

The solution to the quadratic for $C$ is,

$$C = \frac{-b \pm \sqrt{b^2 - 4ac}}{2a} \qquad \text{Eq. 3.14}$$

*Where,*

$a = \beta^2$

$b = -\beta(R_{tot} + S_{tot}) - \frac{\gamma}{\boldsymbol{\alpha}}$

$c = S_{tot}R_{tot}$

Hence, a final expression for computing $C$ is given by (Eq. 3.15),

$$C = \frac{-\left(-\beta(R_{tot} + S_{tot}) - \frac{\gamma}{\boldsymbol{\alpha}}\right) \pm \sqrt{\left(-\beta(R_{tot} + S_{tot}) - \frac{\gamma}{\boldsymbol{\alpha}}\right)^2 - 4\beta^2(S_{tot}R_{tot})}}{2\beta^2}$$

The above expression describes the concentration of the bound riboswitch with pseudoknot as a function of the total concentrations of reactants, $R_{tot}$ and $S_{tot}$ and the equilibrium constants.

***Determination of Model 3 parameters***

Assuming this model, the overall (apparent) dissociation constant using Isothermal calorimetry (ITC) which was reported by Gilbert *et al.* (3) ($Kd_{ITC} = 0.67 \pm 0.06\ \mu M$) would describe the following reaction,

$$Unbound\ RNA + SAM \underset{Ka_{ITC}}{\Longleftrightarrow} bound\ RNA$$

$$\left[R_0 + R + R_{ter} + R_{terMg}\right] + [S] \underset{Ka_{ITC}}{\Longleftrightarrow} [I + C]$$

$$[R_0 + R + R_{terE}] + [S] \underset{Ka_{ITC}}{\Longleftrightarrow} [I + C]$$

Hence, this experimental dissociation constant describes the following equilibrium by,

$$Ka_{ITC} = \frac{Bound\ RNA}{Unbound\ RNA\ \times free\ SAM}$$

$$= \frac{[I + C]}{\left[R_0 + R + R_{ter} + R_{terMg}\right] \times [S]}$$

$$= \frac{[I + C]}{[R_0 + R + R_{terE}] \times [S]}$$

$$= \frac{\left[\left(\frac{1}{K_{fold}.Mg}\right).C + C\right]}{\left[\left(\frac{1}{K_{conv1}K_{terE}.Mg}\right)R_{terE} + \left(\frac{1}{K_{terE}.Mg}\right)R_{terE} + R_{terE}\right] \times [S]}$$

$$= \frac{C\left(\frac{1}{K_{fold}.Mg} + 1\right)}{R_{terE} \times S \times \left(\frac{1}{K_{conv1}K_{terE}.Mg} + \frac{1}{K_{terE}.Mg} + 1\right)}$$

$$= \frac{C \times \beta_1}{R_{terE} \times S \times \alpha_1}$$

Thus,

$$Ka_{ITC} = K_{bindE}\ \frac{\beta_1}{\alpha_1} \qquad \text{Eq. 3.16}$$

*where,* $K_{bindE}$ is the SAM binding association constant of $R_{terE}$, $\beta_1 = \left(\frac{1}{K_{fold}\,.\,Mg} + 1\right)$ and , $\alpha_1 = \left(\frac{1}{K_{conv1}\ K_{terE}\,.\,Mg} + \frac{1}{K_{terE}\,.\,Mg} + 1\right)$. The value of $\alpha_1$ is given, and depends on the $Mg^{2+}$ ions concentration, whereas $\beta_1$ is unknown and cannot be determined from the thermodynamic cycle alone.

Similarly,

$$Ka_{ITC} = \frac{Bound\ RNA}{Unbound\ RNA\ \times free\ SAM}$$

$$= \frac{[I + C]}{\left[R_0 + R + R_{ter} +\ R_{terMg}\right] \times [S]}$$

$$= \frac{[I + C]}{[R_0 + R + R_{terE}] \times [S]}$$

$$= \frac{\left[I + K_{fold}.\ I\ .Mg\right]}{\left[\frac{1}{K_{conv1}}R + R + K_{terE}\ .Mg\ .\ R\right] \times [S]}$$

$$= \frac{I\left(K_{fold}.Mg + 1\right)}{R \times S \times \left(\frac{1}{K_{conv1}} + K_{terE}\ .Mg + 1\right)}$$

$$= \frac{I \times \beta_2}{R \times S \times \alpha_2}$$

Thus,

$$Ka_{ITC} =\ K_{bind}\ \frac{\beta_2}{\alpha_2} \qquad \text{Eq. 3.17}$$

*where,* $K_{bind}$ is the SAM binding association constant of $R$, $\alpha_2 = \left(\frac{1}{K_{conv1}} + K_{terE}\ .Mg + 1\right)$ and $\beta_2 = \left(K_{fold}\ .Mg + 1\right)$.

As in previous models, we consider that the previously reported FS studies monitored the RNA folding in the presence of the ligand relative to its absence (4). In other words, the apparent binding constant determines the fraction of RNA species with a fully formed pseudoknot in the presence of the ligand relative to the ones with partially formed pseudoknot in the absence of the ligand. Therefore, the binding constants deduced from these FS studies, represent the following reaction,

$$Folded\ \&\ unbound\ RNA + SAM \xleftrightarrow[Ka_{FS}]{} Folded\ \&\ bound\ RNA$$

$$\left[R_{ter} +\ R_{terMg}\right] + [S]\ \xleftrightarrow[Ka_{FS}]{}\ [C]$$

$$[R_{terE}] + [S] \underset{Ka_{FS}}{\Longleftrightarrow} [C]$$

$$Ka_{FS} = \frac{Bound\ RNA\ species\ with\ a\ pseudoknot}{Unbound\ RNA\ species\ with\ a\ pseudoknot\ \ \times free\ SAM}$$

$$= \frac{[C]}{[R_{ter} + \ R_{terMg}] \times [S]} = \frac{[C]}{[R_{terE}] \times [S]} = \ K_{bindE}$$

Eq. 3.18

From Eq. 3.16 and Eq. 3.18, it can be observed that,

$$Ka_{ITC} \quad = Ka_{FS}\ \frac{\beta_1}{\alpha_1}$$

$$\beta_1 = \frac{Ka_{ITC}\ \alpha_1}{Ka_{FS}} = \ 1 + \frac{1}{K_{fold}\ .\ Mg}$$

Thus,

$$Kd_{fold} = \left[\frac{Ka_{ITC}\ \alpha_1}{Ka_{FS}} - 1\right] Mg$$

$$= \left[\frac{Ka_{ITC}\ \alpha_1}{Ka_{FS}} - 1\right] Mg$$

$Where,\ \alpha_1 = \left(\frac{1}{K_{conv1}\ K_{terE}\ .\ Mg} + \frac{1}{K_{terE}\ .\ Mg} + 1\right).$ Eq. 3.19

Based on the $Mg^{2+}$ binding model (1) we can therefore determine $K_{terE}$ as follows,

$$K_{terE} = \frac{R_{terE}}{R \times Mg}$$

$$= \frac{R_{ter} + R_{terMg}}{R \times Mg}$$

$$= \frac{K_{conv2}.R + K_{conv2}\ .R\ .Mg\ .K_{Mg}}{R \times Mg}$$

$$= K_{conv2}\frac{1}{Mg} + K_{conv2}.K_{Mg}$$

$$= \ K_{conv2}\left(\frac{1}{Mg} + K_{Mg}\right)$$

Eq. 3.20

Since $K_{conv2}$ and $K_{Mg}$ are known, $K_{terE}$ can be calculated. Consequently, $Kd_{fold}$ for this model can be computed based on $Ka_{ITC}$ and $Ka_{FS}$. $Kd_{fold}$ can also be obtained with the aid of the thermodynamic cycle shown in **Figure 2B**). As in all models, the calculated values for the equilibrium constants are based on the model assumptions.

### 1.5 *Model 4*

This model (illustrated in **Figure 1D**) is similar to the *Mg$^{2+}$ Model* (two-step single ligand) previously described in ref (1), except the ligand here is SAM and the final conformer is denoted as $I$. For convenance, the summary and final equations are reviewed below. From the mass conservation law and **Figure 1D**, the following can be defined,

$$K_{ter} = \frac{R_{ter}}{R^{`}} \qquad \text{Eq. 4.1}$$

$$K_{bindter} = \frac{I}{R_{ter} \times S} \qquad \text{Eq. 4.2}$$

$$I = K_{bindter} \times R_{ter} \times S = K_{bindter} K_{ter}.R^{`}.S \qquad \text{Eq. 4.3}$$

Taking into account that, $S_{tot} = S + I$ and $R_{tot} = R^{`} + R_{ter} + I$ (by substituting in equation 4.3), and solving for the species $I$ will lead to a quadratic that is given by Eq. 4.4,

$$I = \frac{S_{tot} + R_{tot} + \frac{1}{\gamma} \pm \sqrt{\left(-S_{tot} - R_{tot} - \frac{1}{\boldsymbol{\alpha}}\right)^2 - 4R_{tot}S_{tot}}}{2}$$

and $\boldsymbol{\alpha} = \frac{K_{bindter}}{\frac{1}{K_{ter}} + 1}$

The quadratic equation has two possible solutions for $I$. Only one of the two roots is physically sensible, and can represent concentrations (i.e. positive real number, with a maximum value equal or less than $R_{tot}$).

### 1.6 *Model 5*

From the definition of equilibrium and **Figure 1E**, the following can be defined,

$$K_{conv1} = \frac{R}{R_0} \quad \text{Eq. 5.1}$$

$$K_{terE} = \frac{R_{terE}}{R} \quad \text{Eq. 5.2}$$

$$K_{bind} = \frac{I}{R \times S} \quad \text{Eq. 5.3}$$

$$K_{bindE} = \frac{C}{R_{terE} \times S} \quad \text{Eq. 5.4}$$

$$K_{fold} = \frac{C}{I} = \frac{R_{terE}\ S\ K_{bindE}}{R\ S\ K_{bind}} = \frac{R\ S\ K_{bindE}\ K_{terE}}{R\ S\ K_{bind}} = \frac{K_{bindE}\ K_{terE}}{K_{bind}} \quad \text{Eq. 5.5}$$

From the definition of equilibrium we have,

$$C = I \times K_{fold}$$

$$= R \times S \times K_{bind} \times K_{fold}$$

By substituting for $K_{fold}$, from equation 5.5, we get

$$= R_0 \times S \times \left( K_{bind} \times K_{conv1} \times \left( \frac{K_{bindE}\ K_{terE}}{K_{bind}} \right) \right)$$

$$= R_0 \times S \times (K_{conv1} \times K_{bindE} \times K_{terE})$$

$$C = R_0 \times S \times \boldsymbol{\alpha} \quad \text{Eq. 5.6}$$

*where*, $\boldsymbol{\alpha} = (K_{conv1} \times K_{bindE} \times K_{terE})$

From the conservation of moieties, the total RNA is equal to,

$$R_{tot} = R_0 + R + R_{terE} + I + C \quad \text{Eq. 5.7}$$

By substitution of equations 5.1, 5.2, 5.5, in equation 5.7,

$$R_{tot} = R_0 + R_0 K_{conv1} + R K_{terE} + \frac{C}{K_{fold}} + C$$

$$= R_0 + R_0K_{conv1} + R_0K_{conv1}K_{terE} + C\left(\frac{1}{K_{fold}} + 1\right)$$

$$R_{tot} = R_0 + R_0K_{conv1} + R_0K_{conv1}K_{terE} + C\beta \qquad \text{Eq. 5.8}$$

*where,*

$$\beta = \left(\frac{1}{K_{fold}} + 1\right) = \frac{K_{bind}}{K_{bindE}\,K_{terE}} + 1 \qquad \text{Eq. 5.9}$$

Thus,

$$R_0 + R_0K_{conv1} + R_0K_{conv1}K_{terE} = R_{tot} - C\beta$$

$$R_0 = \frac{R_{tot} - C\beta}{1 + K_{conv1} + K_{conv1}K_{terE}} = \frac{R_{tot} - C\beta}{\gamma} \qquad \text{Eq. 5.10}$$

And

$$\gamma = 1 + K_{conv1} + K_{conv1}\,K_{terE} = 1 + K_{conv1}(1 + K_{terE}) \qquad \text{Eq. 5.11}$$

The total SAM concentration $S_{tot}$ , is a conserved quantity and is equal to

$$S_{tot} = S + I + C \qquad \text{Eq. 5.12}$$

Thus, the free SAM is described by,

$$S = S_{tot} - (I + C)$$

$$= S_{tot} - \left(\frac{C}{K_{fold}} + C\right)$$

$$= S_{tot} - C\left(\frac{1}{K_{fold}} + 1\right)$$

$$S = S_{tot} - C\beta \qquad \text{Eq. 5.13}$$

To obtain the concentrations of the bound aptamer, $C$, in terms of the equilibrium constants and the total concentration of the reactants, $R_{tot}$ and $S_{tot}$ , we substitute for $R_0$ and $S$ from equations 5.10 and 5.13 in equation 5.6,

$$C = R_0 \times S \times \boldsymbol{\alpha}$$

$$= \left(\frac{R_{tot} - C\beta}{\gamma}\right) \times (S_{tot} - C\beta) \times \boldsymbol{\alpha}$$

$$= (R_{tot} - C\beta) \times (S_{tot} - C\beta) \times \frac{\boldsymbol{\alpha}}{\gamma}$$

Thus, $\frac{\gamma}{\boldsymbol{\alpha}} C = (R_{tot} - C\beta) \times (S_{tot} - C\beta)$

By distributing the brackets, subtracting $\frac{\gamma}{\boldsymbol{\alpha}} C$ from both sides and taking $C$ as a common factor, we get a quadratic,

$$\beta^2 C^2 - R_{tot}\beta C - S_{tot}\beta C - \frac{\gamma}{\boldsymbol{\alpha}} C + S_{tot}R_{tot} = 0$$

$$(\beta^2)C^2 + \left(-\beta(R_{tot} + S_{tot}) - \frac{\gamma}{\boldsymbol{\alpha}}\right) C + (S_{tot}R_{tot}) = 0$$

The solution to the quadratic for $C$ is,

$$C = \frac{-b \pm \sqrt{b^2 - 4ac}}{2a} \qquad \text{Eq. 5.14}$$

*Where,*

$$a = \beta^2$$

$$b = -\beta(R_{tot} + S_{tot}) - \frac{\gamma}{\boldsymbol{\alpha}}$$

$$c = S_{tot}R_{tot}$$

Hence, a final expression for computing $C$ is given by (Eq. 5.15),

$$C = \frac{-\left(-\beta(R_{tot} + S_{tot}) - \frac{\gamma}{\boldsymbol{\alpha}}\right) \pm \sqrt{\left(-\beta(R_{tot} + S_{tot}) - \frac{\gamma}{\boldsymbol{\alpha}}\right)^2 - 4\beta^2(S_{tot}R_{tot})}}{2\beta^2}$$

The above expression describes the concentration of the bound riboswitch with pseudoknot as a function of the total concentrations of reactants, $R_{tot}$, $Mg$ and $S_{tot}$ and the equilibrium constants.

***Determination of Model 5 parameters***

To be able to apply this model, the equilibrium constants $K_{conv1}$ , $K_{terE}$, $K_{bind}$, $K_{bindE}$ and $K_{fold}$ needs to be estimated from existing experimental values. The equilibrium constant $K_{conv1}$ can be obtained from the calculated binding energies and from the probabilities of the binding *competent* species ($R$) and binding *incompetent* ($R_0$) species. The model considers the effect of differential binding of riboswitch with pseudoknot ($R_{terE}$) and intermediate which is the riboswitch without a pseudoknot ($R$) into account. Since $Mg^{2+}$ concentration is assumed negligible (or with constant effect), $K_{terE}$ can be calculated as the ratio between $\frac{R_{terE}}{R}$.

To determine $K_{terE}$ at a given fixed magnesium concentration it is useful to not that,

$$K_{terE} = \frac{RNA\ with\ the\ accessible\ binding\ contacts\ and\ with\ pseudoknot\ in\ the\ presence\ of\ Mg}{RNA\ with\ the\ accessible\ binding\ contacts\ and\ without\ pseudoknot\ in\ the\ presence\ of\ Mg}$$

$$= \frac{fraction\ binding\ competent\ at\ a\ given\ Mg\ concentration}{R}$$

$$= \frac{R_{terE}}{R} = \frac{R_{ter} + R_{terMg}}{R} = \frac{R_{ter}}{R} + \frac{R_{terMg}}{R} = K_{conv2} + \left(K_{conv2} \times \frac{R_{terMg}}{R_{ter}}\right)$$

*where*, $K_{conv2}$ and $\frac{R_{terMg}}{R_{ter}}$ can be computed based on $Mg^{2+}$ data and model, as shown in our initial study (1), at specific $Mg^{2+}$ concentrations (the total $Mg^{2+}$ concentration is assumed to be 2 mM). The accessible contacts in the above equation refer to the population with the correct secondary structure. This assumes that the riboswitch will not be affected by the magnesium concentration (i.e. $Mg^{2+}$ concentration has no effect on the $R_{terE}$). The consequence of the latter assumption is that the fraction having a pseudoknot in the absence of SAM is fixed and generally independent of the $Mg^{2+}$ concentration. The latter approximation/assumption may become valid if, for example, the consumption of $R_{terE}$ is very limited i.e. if the binding of $R_{terE}$ is very weak.

To ease the link between the model parameters and the measured (apparent) experimental data, we define $K_2$ be the equilibrium determined from experimentally measured probabilities (presence and absence of pseudoknot), in the presence of a fixed $Mg^{2+}$ concentration. Thus, from the definition of equilibrium of $K_{terE}$ (Eq. 5.2, and illustrated in **Figure 1E**) we have,

$$K_2 \quad = \frac{RNA\ with\ pseudoknot}{RNA\ without\ pseudoknot}$$

$$= \frac{R_{terE}}{R^{`}} = \frac{R_{terE}}{R_0 + R} = \frac{R_{terE}}{R\left(\frac{1}{K_{conv1}} + 1\right)} = K_{terE}\frac{1}{\left(\frac{1}{K_{conv1}} + 1\right)} \qquad \text{Eq. 5.16}$$

Like in previous models, we make use of the fact that the overall (apparent) dissociation constant was determined experimentally using Isothermal calorimetry (ITC), reported by Gilbert *et al.* (3), $Kd_{ITC}$ was experimentally determined to be $0.67 \pm 0.06\ \mu M$.

Assuming this model, the latter apparent reaction equilibrium constant would describe the following reaction,

$$Unbound\ RNA + SAM \xLeftrightarrow[Ka_{ITC}]{} bound\ RNA$$

$$[R_0 + R + R_{terE}] + [S] \xLeftrightarrow[Ka_{ITC}]{} [I + C]$$

$$\left[R^{`} + R_{terE}\right] + [S] \xLeftrightarrow[Ka_{ITC}]{} [I + C]$$

Hence, this experimental dissociation constant describes the following equilibrium by,

$$Ka_{ITC} = \frac{Bound\ RNA}{Unbound\ RNA\ \times free\ SAM}$$

$$= \frac{[I + C]}{[R_0 + R + R_{terE}] \times [S]}$$

$$= \frac{[I + C]}{[R^{`} + R_{terE}] \times [S]}$$

$$= \frac{\left[\left(\frac{1}{K_{fold}}\right).C + C\right]}{\left[\left(\frac{1}{K_2}\right).R_{terE} + R_{terE}\right] \times [S]}$$

$$= \frac{C\left({}^{1}/_{K_{fold}} + 1\right)}{R_{terE} \times S\left({}^{1}/_{K_2} + 1\right)}$$

$$= \frac{C \times \beta_1}{R_{terE} \times S \times \alpha_1}$$

Thus,

$$Ka_{ITC} \quad = K_{bindE} \ \frac{\beta_1}{\alpha_1} \qquad \text{Eq. 5.17}$$

*where,* $K_{bindE}$ is the SAM binding association constant of $R_{terE}$ , $\alpha_1 = \left(1/_{K_2} + 1\right)$ and $\beta_1 = \left(1/_{K_{fold}} + 1\right)$. It is worth noting that $\alpha_1$ can be re-written in terms of $K_{terE}$ and $K_{conv1}$ by substituting for $K_2$ (Eq. 5.16) to yield in $\alpha_1 = \frac{1}{K_{terE}\,K_{conv1}} + K_{terE} + 1$. The value of $K_{conv1}$is readily available from base pairing probabilities and $K_{terE}$ can be computed at a specific $Mg^{2+}$ concentration. Thus, the value of $\alpha_1$ can be obtained at an assumed $Mg^{2+}$ concentration, whereas $\beta_1$ is unknown and cannot be determined from the thermodynamic cycle alone.

Similarly (and alternatively),

$$Ka_{ITC} = \frac{Bound\ RNA}{Unbound\ RNA\ \times free\ SAM}$$

$$= \frac{[I + C]}{[R_0 + R + R_{terE}] \times [S]}$$

$$= \frac{[I + C]}{[R` + R_{terE}] \times [S]}$$

$$= \frac{\left[I + K_{fold}.I\right]}{[R + K_{terE}.R] \times [S]}$$

$$= \frac{I\left(1 + K_{fold}\right)}{R \times S(1 + K_{terE})}$$

$$= \frac{I \times \beta_2}{R \times S \times \alpha_2}$$

Thus,

$$K_{ITC} = \ K_{bind} \ \frac{\beta_2}{\alpha_2} \qquad \text{Eq. 5.18}$$

*where,* $K_{bind}$ is the SAM binding association constant of $R$, $\alpha_2 = (1 + K_{terE})$ and $\beta_2 = \left(1 + K_{fold}\right)$.

It should be noted that, the value of the parameter $K_{bindE}$ in this model is considered to be equivalent to the constant that has been reported experimentally by Haller *et al* (4), that is we have $K_{bindE} \approx K_{FS}$. The latter was used in subsequent calculations. Thus, from the above expression for $Ka_{ITC}$ , $K_{fold}$ is given by,

$$K_{fold} = \frac{1}{\frac{Kd_{bindE} \, . \, \alpha_1}{Kd_{ITC}} - 1} = \frac{1}{\frac{Kd_{FS} \, . \, \alpha_1}{Kd_{ITC}} - 1} \qquad \text{Eq. 5.19}$$

Hence, $Kd_{fold} = \frac{1}{K_{fold}}$ and $dG_{fold} = -RT \ln\left(K_{fold}\right)$ can be obtained from Eq. 5.19. Since three of the four parameters are known, then the fourth can be determined through a thermodynamic cycle.

Hence, we construct a thermodynamic cycle (**Figure 2C**), *where,*

$$\Delta G_{bind} + \Delta G_{fold} + (-\Delta G_{bindE}) + (-\Delta G_{Conv2}) = 0$$

By rearrangement, $dG_{bind}$ can be computed from,

$$\Delta G_{bind} = -\Delta G_{fold} - (-\Delta G_{bindE}) - (-\Delta G_{Conv2})$$

Eq. 5.20

### 1.7 *Model 6*

From the definition of equilibrium and **Figure 1F**, the following can be defined,

$$K_{conv1} = \frac{R}{R_0} \qquad \text{Eq. 6.1}$$

$$K_{conv2} = \frac{R_{ter}}{R} \qquad \text{Eq. 6.2}$$

$$K_{bind} = \frac{I}{R \times S} \qquad \text{Eq. 6.3}$$

$$K_{bindter} = \frac{C}{R_{ter} \times S} \qquad \text{Eq. 6.4}$$

$$K_{fold} = \frac{C}{I} = \frac{R_{ter}\ S\ K_{bindter}}{R\ S\ K_{bind}} = \frac{R\ S\ K_{bindter}\ K_{conv2}}{R\ S\ K_{bind}} = \frac{K_{bindter}\ K_{conv2}}{K_{bind}} \qquad \text{Eq. 6.5}$$

From the definition of equilibrium (in eq. 6.5) we have,

$$C = I \times K_{fold}$$

$$= R \times S \times K_{bind} \times K_{fold}$$

By substituting for $K_{fold}$, from equation 6.5, we get

$$= R_0 \times S \times \left( K_{bind} \times K_{conv1} \times \left( \frac{K_{bindter}\ K_{conv2}}{K_{bind}} \right) \right)$$

$$= R_0 \times S \times (K_{conv1} \times K_{bindter} \times K_{conv2})$$

$$C = R_0 \times S \times \boldsymbol{\alpha} \qquad \text{Eq. 6.6}$$

*where*, $\boldsymbol{\alpha} = (K_{conv1} \times K_{bindter} \times K_{conv2})$

From the conservation of moieties, the total RNA is equal to,

$$R_{tot} = R_0 + R + R_{ter} + I + C \qquad \text{Eq. 6.7}$$

By substitution of equations 6.1, 6.2, 6.5, in equation 6.7,

$$R_{tot} \quad = \; R_0 + \; R_0K_{conv1} + RK_{conv2} + \frac{C}{K_{fold}} + C$$

$$= \; R_0 + \; R_0K_{conv1} + R_0K_{conv1}K_{conv2} + C\left(\frac{1}{K_{fold}} + 1\right)$$

$$R_{tot} \quad = \; R_0 + \; R_0K_{conv1} + R_0K_{conv1}K_{conv2} + C\beta \qquad \text{Eq. 6.8}$$

*where,*

$$\beta = \left(\frac{1}{K_{fold}} + 1\right) = \; \frac{K_{bind}}{K_{bindter}\,K_{conv2}} + 1 \qquad \text{Eq. 6.9}$$

Thus,

$$R_0 + R_0K_{conv1} + R_0K_{conv1}K_{conv2} \; = \; R_{tot} - \; C\beta$$

$$R_0 = \frac{R_{tot} - \; C\beta}{1 + K_{conv1} + K_{conv1}K_{conv2}} = \; \frac{R_{tot} - \; C\beta}{\gamma} \qquad \text{Eq. 6.10}$$

And

$$\gamma = 1 + K_{conv1} + K_{conv1}\,K_{conv2} = 1 + K_{conv1}(1 + K_{conv2}) \qquad \text{Eq. 6.11}$$

The total SAM concentration $S_{tot}$ , is a conserved quantity and is equal to

$$S_{tot} = S + I + C \qquad \text{Eq. 6.12}$$

Thus, the free SAM is described by,

$$S = S_{tot} - (I + C)$$

$$= S_{tot} - \left(\frac{C}{K_{fold}} + C\right)$$

$$= S_{tot} - C\left(\frac{1}{K_{fold}} + 1\right)$$

$$S = \; S_{tot} - C\beta \qquad \text{Eq. 6.13}$$

To obtain the concentrations of the bound aptamer, $C$, in terms of the equilibrium constants and the total concentration of the reactants, $R_{tot}$ and $S_{tot}$ , we substitute for $R_0$ and $S$ from equations 6.10 and 6.13 in equation 6.6,

$$C = R_0 \times S \times \boldsymbol{\alpha}$$

$$= \left(\frac{R_{tot} - C\beta}{\gamma}\right) \times (S_{tot} - C\beta) \times \boldsymbol{\alpha}$$

$$= (R_{tot} - C\beta) \times (S_{tot} - C\beta) \times \frac{\boldsymbol{\alpha}}{\gamma}$$

Thus,

$$\frac{\gamma}{\boldsymbol{\alpha}}C = (R_{tot} - C\beta) \times (S_{tot} - C\beta)$$

By distributing the brackets, subtracting $\frac{\gamma}{\boldsymbol{\alpha}}C$ from both sides and taking $C$ as a common factor, we get a quadratic,

$$\beta^2C^2 - R_{tot}\beta C - S_{tot}\beta C - \frac{\gamma}{\boldsymbol{\alpha}}C + S_{tot}R_{tot} = 0$$

$$(\beta^2)C^2 + \left(-\beta(R_{tot} + S_{tot}) - \frac{\gamma}{\boldsymbol{\alpha}}\right)C + (S_{tot}R_{tot}) = 0$$

The solution to the quadratic for $C$ is,

$$C = \frac{-b \pm \sqrt{b^2 - 4ac}}{2a} \qquad \text{Eq. 6.14}$$

*Where,*

$$a = \beta^2$$

$$b = -\beta(R_{tot} + S_{tot}) - \frac{\gamma}{\boldsymbol{\alpha}}$$

$$c = S_{tot}R_{tot}$$

Final equation, Eq. 6.15:

$$C = \frac{-\left(-\beta(R_{tot} + S_{tot}) - \frac{\gamma}{\boldsymbol{\alpha}}\right) \pm \sqrt{\left(-\beta(R_{tot} + S_{tot}) - \frac{\gamma}{\boldsymbol{\alpha}}\right)^2 - 4\beta^2(S_{tot}R_{tot})}}{2\beta^2}$$

The above expression describes the concentration of the bound riboswitch with pseudoknot as a function of the total concentrations of reactants, $R_{tot}$, $Mg$ and $S_{tot}$ and the equilibrium constants.

***Determination of Model 6 parameters***

Again, as in other models, that fact that the overall (apparent) dissociation constant had been determined experimentally using Isothermal calorimetry (ITC), and was reported by Gilbert *et al* (3) is used to determine the model parameters. In the latter study, the authors reported a dissociation constant of $Kd_{ITC}$ of $0.67 \pm 0.06\ \mu M$.

If this model, *Model 6*, was true, then the latter apparent reaction association equilibrium constant ($Ka_{ITC} = \frac{1}{Kd_{ITC}}$) would describe the following reaction,

$$Unbound\ RNA + SAM \underset{Ka_{ITC}}{\Longleftrightarrow} bound\ RNA$$

$$[R_0 + R + R_{ter}] + [S] \underset{Ka_{ITC}}{\Longleftrightarrow} [I + C]$$

$$\left[R^{`} + R_{ter}\right] + [S] \underset{Ka_{ITC}}{\Longleftrightarrow} [I + C]$$

Hence, the experimental association constant $Ka_{ITC}$ would be then described by the following equilibrium,

$$Ka_{ITC} = \frac{Bound\ RNA}{Unbound\ RNA\ \times free\ SAM}$$

$$= \frac{[I + C]}{[R_0 + R + R_{ter}] \times [S]}$$

$$= \frac{[I + C]}{[R^{`} + R_{ter}] \times [S]}$$

By substituting for the equilibrium constants, as defined in the in the model derivation, Equations 6.1 to 6.5, we therefore obtain,

$$= \frac{\left[\left(\frac{1}{K_{fold}}\right).C + C\right]}{\left[(\frac{1}{K_{ter}}).R_{ter} + R_{ter}\right] \times [S]}$$

$$= \frac{C\left(1/_{K_{fold}} + 1\right)}{R_{ter} \times S\left(1/_{K_{ter}} + 1\right)}$$

$$= \frac{C \times \beta_1}{R_{ter} \times S \times \alpha_1}$$

Thus,

$$Ka_{ITC} \quad = K_{bindter} \ \frac{\beta_1}{\alpha_1} \qquad \text{Eq. 6.16}$$

*where,* $K_{bindter}$ is the SAM binding association constant of $R_{ter}$ , $\alpha_1 = \left({}^{1}/_{K_{ter}} + 1\right)$ and $\beta_1 = \left({}^{1}/_{K_{fold}} + 1\right)$. The value of $\alpha_1$ is given, and according to the $Mg^{2+}$ ions concentration, whereas $\beta_1$ is unknown and cannot be determined from the thermodynamic cycle alone.

Similarly (and alternatively),

$$Ka_{ITC} = \frac{Bound\ RNA}{Unbound\ RNA\ \times free\ SAM}$$

$$= \frac{[I + C]}{[R_0 + R + R_{ter}] \times [S]}$$

$$= \frac{\left[I + K_{fold}.I\right]}{[(1/K_{conv1})R + R + K_{conv2}.R] \times [S]}$$

$$= \frac{I\left(K_{fold} + 1\right)}{R \times S\left((1/K_{conv1}) + K_{conv2} + 1\right)}$$

$$= \frac{I \times \beta_2}{R \times S \times \alpha_2}$$

Thus,

$$Ka_{ITC} = \ K_{bind} \ \frac{\beta_2}{\alpha_2} \qquad \text{Eq. 6.17}$$

*where,* $K_{bind}$ is the SAM binding association constant of $R$, $\alpha_2 = \left((1/K_{conv1}) + K_{conv2} + 1\right)$ and $\beta_2 = \left(K_{fold} + 1\right)$.

As in the previous model, the value of $K_{bindter} \approx K_{FS}$ is a parameter that has been reported experimentally by Haller *et al* (4), which was used in subsequent calculations. Hence, based on the above expression for $Ka_{ITC}$ , $K_{fold}$ is given by,

$$K_{fold} = \frac{1}{\frac{Kd_{bindter}\,.\,\alpha_1}{Kd_{ITC}} - 1} = \frac{1}{\frac{Kd_{FS}\,.\,\alpha_1}{Kd_{ITC}} - 1} \quad \text{Eq. 6.18}$$

Hence, $Kd_{fold} = \frac{1}{K_{fold}}$ and $dG_{fold} = -RT\ln(K_{fold})$ can be computed from Eq. 6.18. Since three $(\Delta G_{fold}, \Delta G_{bindter}\ and\ \Delta G_{Conv2})$ of the four parameters are known, then the fourth can be determined through a thermodynamic cycle.

Hence, we construct a thermodynamic cycle (**Figure 2D**), *where,*

$$\Delta G_{bind} + \Delta G_{fold} + (-\Delta G_{bindter}) + (-\Delta G_{Conv2}) = 0$$

By rearrangement, $dG_{bind}$ is given by,

$$\Delta G_{bind} = -\Delta G_{fold} - (-\Delta G_{bindter}) - (-\Delta G_{Conv2})$$

Eq. 6.19

### 2 Supplementary Figures

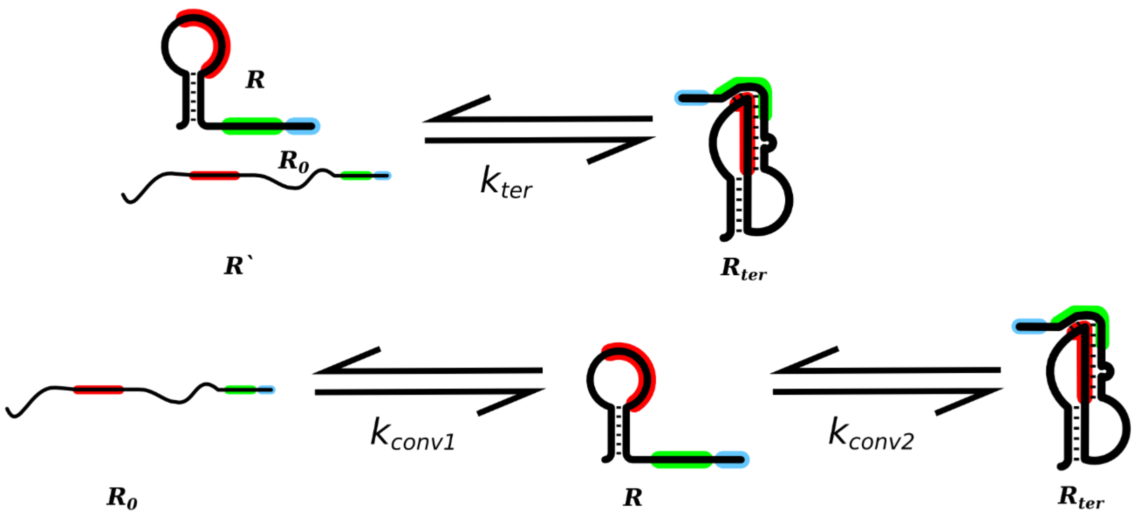


**Figure S1. RNA folding models in the absence of ligands (*Model 0*).** The figure illustrates schemes for RNA folding steps in the absence of any ligands that are used in this study (designated as *Model 0*). For simplicity and to facilitate the link between experimental data and the theoretical models, folding is represented as two equivalent forms, a single step folding scheme (upper panel) or as a two-step folding scheme (lower panel). The related equilibrium constants are shown on the figure. These representations are identical to what have been previously described and used in $Mg^{2+}$ binding models (1).

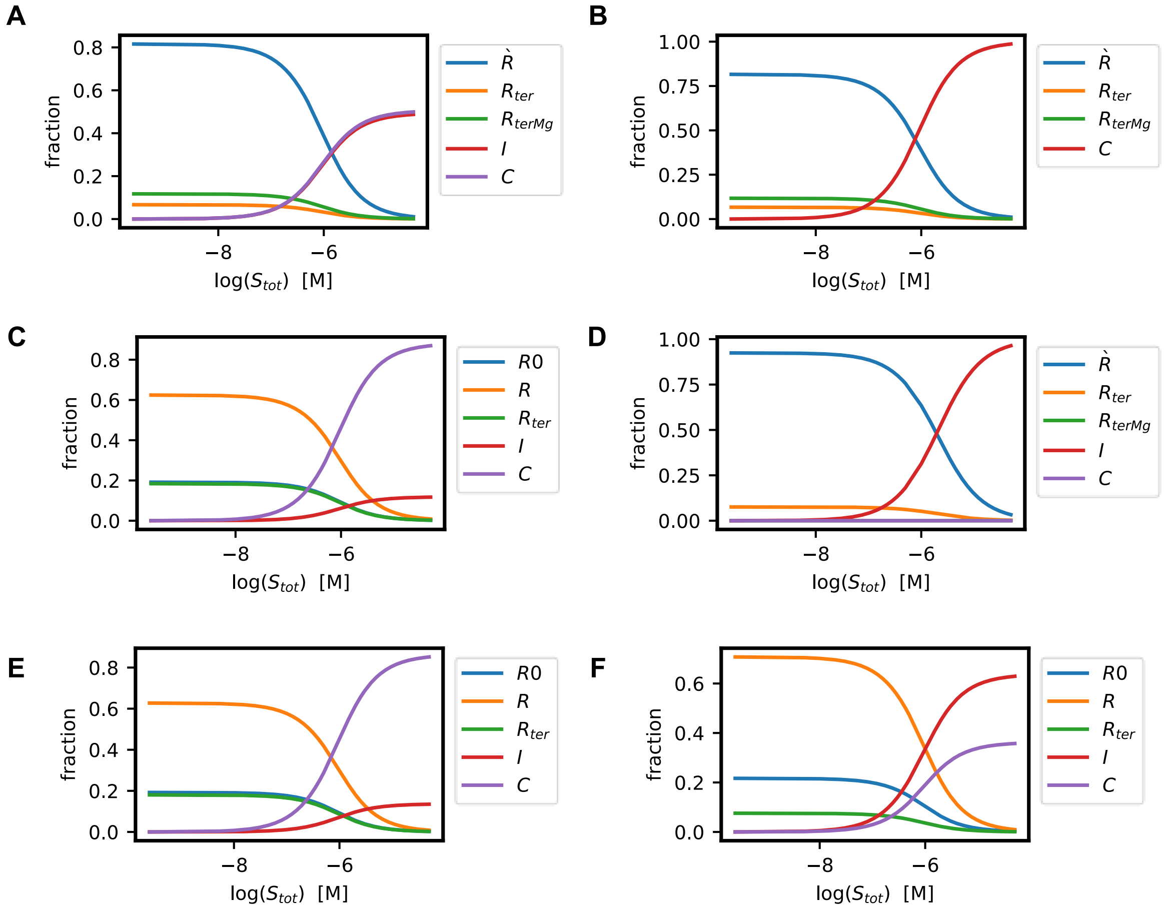


**Figure S2. Predicted titration curves of individual RNA species based on *Models 2-6*.** The Figure shows plots for the fractions of the different RNA species that are computed from *Models 1- 6*, as a function of SAM concentration at a fixed total $Mg^{2+}$ concentration of 2.0 mM. Similar titration curves for individual RNA species computed based on *Model 1* are illustrated in **Figure 4**.

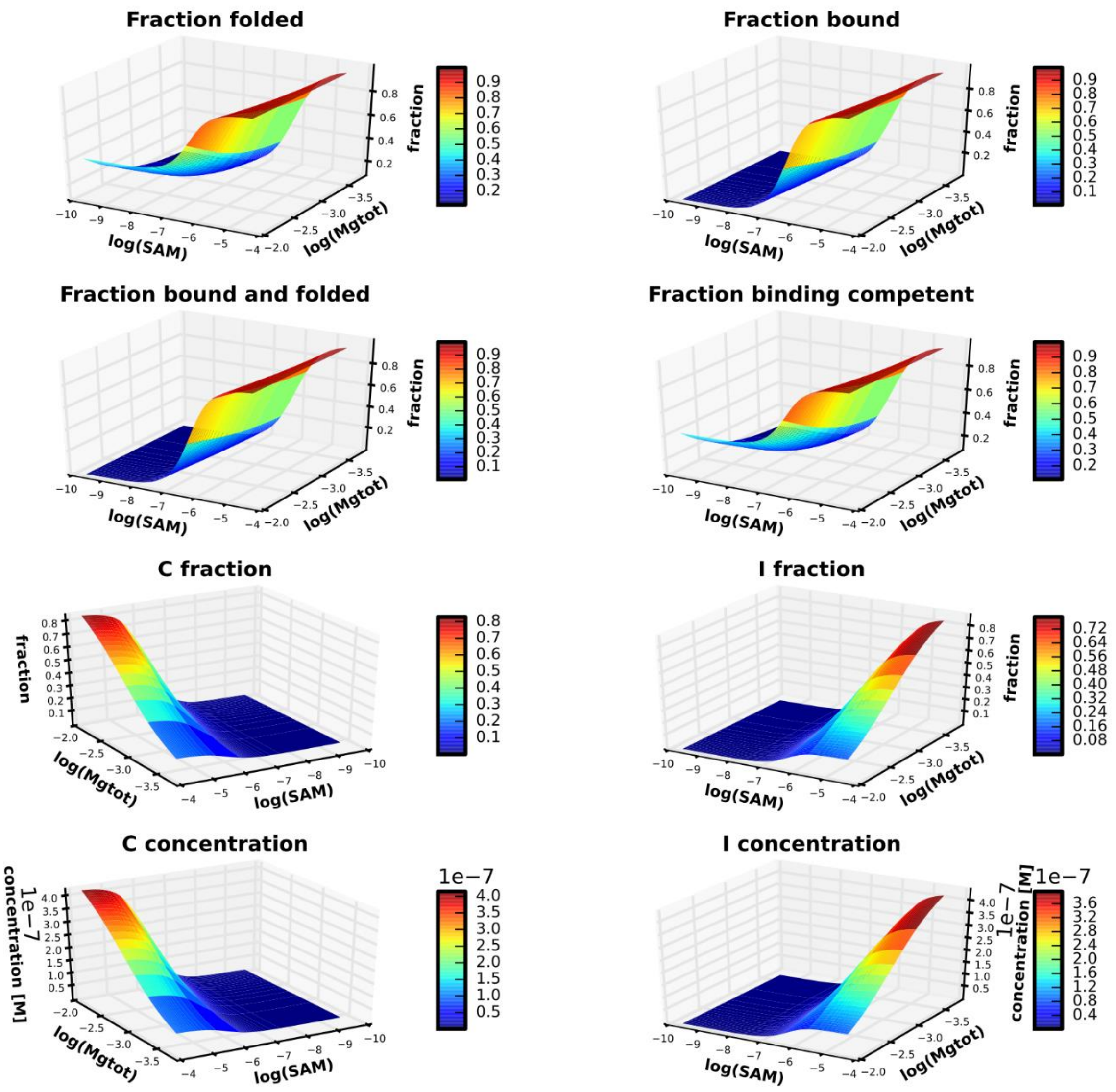


**Figure S3. Concentration dependent titration curves obtained from modeling the SAM-II riboswitch as a function of both SAM and $Mg^{2+}$.** The figure illustrates the fraction forming a pseudoknot (*fraction folded*), *fraction bound*, *fraction bound & folded* and *fraction binding competent* RNA, as well as the fractions (and concentrations) of the conforms $I$ and $C$ of the SAM-II riboswitch, as a function of both the total $Mg^{2+}$ ($Mgtot$) and total SAM concentrations, computed using an RNA concentration of 0.5 μM. The titration curves are computed based on *Model 1* which is illustrated in **Figure 1A** in the main text.

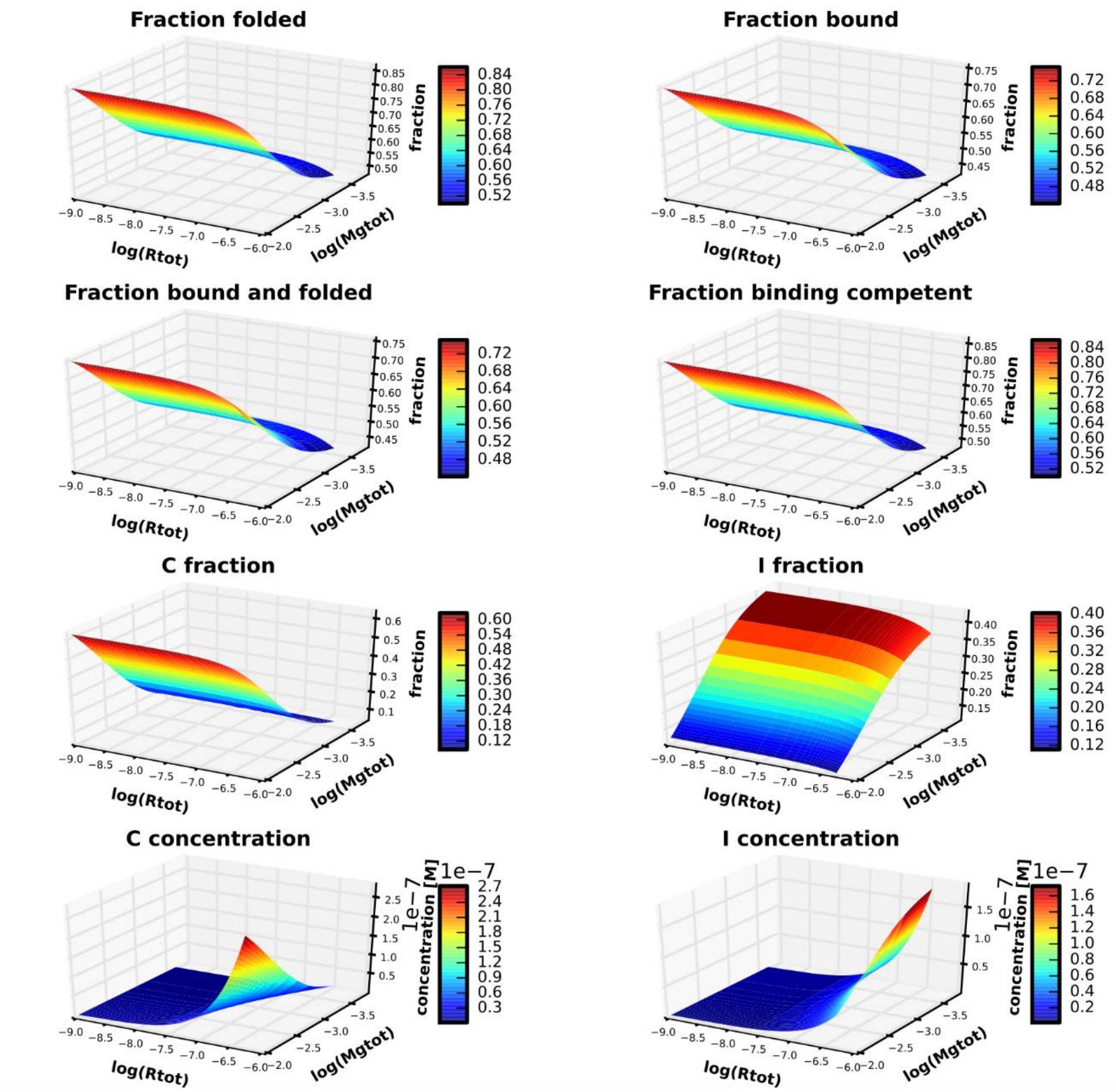


**Figure S4. Three-dimensional concentration dependent titration curves obtained from modeling the SAM-II riboswitch as a function of both $Mg^{2+}$ and RNA at a fixed concentration of SAM.** The figure illustrates the fraction forming a pseudoknot (*fraction folded*), *fraction bound*, the *fraction bound & folded*, and the *fraction binding competent* RNA of the SAM-II riboswitch as a function of both the total $Mg^{2+}$ ($Mgtot$) and total RNA ($Rtot$) concentrations. Further, the fractions (and concentrations) of the bound and folded riboswitch species $I$ and $C$ are also shown. The titration curves are computed based on *Model 1* which is illustrated in **Figure 1A** in the main text, using a total SAM concentration of 1 μM.

### 3 Supplementary Tables

| Model 2 | |
|---|---|
| **Constant** | **Value** |
| Given $K_{Mg}$ from *Mg Model* (1) | 878 |
| Given $K_{ter}$ from *Mg Model* (1) | 0.0819 |
| Given $Kd_{ITC}$ from experiment (3) | 6.70E-07 |
| **Parameters in the presence of $Mg^{2+}$ (kcal/mol):** | |
| $dG_{Mg}$ | -4.016 |
| $dG_{ter}$ | 1.483 |
| $dG_{bindMg}$ | -9.289 |
| $dG_{FS}$ (calculated from model) | -9.425 |
| ***Model 3*** | |
| **Constant** | **Value** |
| Given $K_{conv1}$ from *Mg Model* (1) | 3.263 |
| Given $K_{conv2}$ from *Mg Model* (1) | 0.107 |
| Given $Kd_{ITC}$ from experiment (3) | 6.70E-07 |
| Given $Kd_{FS}$ from experiment (4) | 1.40E-07 |
| $dG_{FS}$ | -9.350 |
| **Parameters in the presence of $Mg^{2+}$(kcal/mol):** | |
| $dG_{conv1}$ | -0.701 |
| $dG_{bindE}$ | -9.350 |
| $dG_{terE}$ | -2.959 |
| $dG_{fold}$ | -4.870 |
| $dG_{bind}$ | -7.439 |
| ***Model 4*** | |
| **Constant** | **Value** |
| Given $K_{ter}$ from *Mg Model* (1) | 0.0819 |
| Given $Kd_{FS}$ from experiment (4) | 1.40E-07 |
| **Parameters in the absence of $Mg^{2+}$(kcal/mol):** | |
| $dG_{ter}$ | 1.483 |
| $dG_{bindter}$ | -9.350 |
| ***Model 5*** | |
| **Constant** | **Value** |
| Given $K_{conv1}$ from *Mg Model* (1) | 3.263 |
| Given $K_{conv2}$ from *Mg Model* (1) | 0.107 |
| Given $Kd_{ITC}$ from experiment (3) | 6.70E-07 |
| Given $Kd_{FS}$ from experiment (4) | 1.40E-07 |
| In the presence of a total $Mg^{2+}$ conc. = 2 mM | |
| $P_{Rter}$ | |
| $P_{RterMg}$ | |

| | |
|---|---|
| Probability of structures with pseudoknot ($P_{Rter}$ + $P_{RterMg}$) | 0.22 |
| Probability of structures without pseudoknot | 0.78 |
| **Parameters in the presence of $Mg^{2+}$ as a constant, in kcal/mol:** | |
| $dG_{terE}$ | 0.7386 |
| $dG_{conv1}$ | -0.7007 |
| $dG_{bindE}$ | -9.350 |
| $dG_{fold}$ | -1.091 |
| $dG_{bind}$ | -7.520 |
| ***Model 6*** | |
| **Constant** | **Value** |
| Given $K_{conv1}$ from *Mg Model* (1) | 3.263 |
| Given $K_{conv2}$ from *Mg Model* (1) | 0.107 |
| Given $Kd_{ITC}$ from experiment (3) | 6.70E-07 |
| Given $Kd_{FS}$ from experiment (4) | 1.40E-07 |
| **Parameters in the absence of $Mg^{2+}$ (kcal/mol):** | |
| $dG_{conv1}$ | -0.701 |
| $dG_{conv2}$ | 1.324 |
| $dG_{bindter}$ | -9.350 |
| $dG_{bind}$ | -8.361 |
| $dG_{fold}$ | 0.335 |

**Table S1.** Equilibrium constants (model parameters) used in the computation of the predicted titration curves for *Models 2 to 6*. The corresponding parameters for *Model 1* are shown in **Table 2** in the main text.

| | $\boldsymbol{\alpha}$ | $\boldsymbol{\beta}$ | $\boldsymbol{\gamma}$ |
|---|---|---|---|
| ***Model 1*** | $K_{ter}.K_{Mg}.Mg.K_{bindMg}$ | $\frac{1}{K_{fold}.Mg}+1$ | $1+K_{ter}+K_{ter}.K_{Mg}.Mg$ |
| ***Model 2*** | $K_{ter}.K_{Mg}.Mg.K_{bindMg}$ | 1.0 | $1+K_{ter}+K_{ter}.K_{Mg}.Mg$ |
| ***Model 3*** | $K_{conv1}.K_{terE}.Mg.K_{bindE}$ | $\frac{1}{K_{fold}.Mg}+1$ | $1+K_{conv1}+K_{conv1}\,K_{terE}.Mg$ |
| ***Model 4*** | $\frac{K_{bindter}}{\frac{1}{K_{ter}}+1}$ | 1.0 | 1.0 |
| ***Model 5*** | $K_{conv1}.K_{terE}.K_{bindE}$ | $\frac{1}{K_{fold}}+1$ | $1+K_{conv1}+K_{conv1}\,K_{terE}$ |
| ***Model 6*** | $K_{conv1}.K_{conv2}.K_{bindter}$ | $\frac{1}{K_{fold}}+1$ | $1+K_{conv1}+K_{conv1}\,K_{conv2}$ |

**Table S2. Summary of Equations for *Models 1-6.*** The solution to the quadratic for concentration of final folded RNA species (depicted as $C$ for all models except for *Model 4* was named *I*) is,

$$C=\frac{-b\pm\sqrt{b^2-4ac}}{2a}$$

*Where,* $a=\beta^2$ $b=-\beta(R_{tot}+S_{tot})-\frac{\gamma}{\alpha}$ $c=S_{tot}R_{tot}$

| Model | $fraction\ bound$ | $fraction\ folded$ | $fraction\ bound\ and\ folded$ | $fraction\ of\ binding\ competent\ RNA$ |
|---|---|---|---|---|
| *Model 1* | $\frac{I+C}{R_{tot}}$ | $\frac{R_{ter}+R_{terMg}+I+C}{R_{tot}}$ | $\frac{I+C}{R_{tot}}$ | $\frac{R_{ter}+R_{terMg}+I+C}{R_{tot}}$ |
| *Model 2* | $\frac{C}{R_{tot}}$ | $\frac{R_{ter}+R_{terMg}+C}{R_{tot}}$ | $\frac{C}{R_{tot}}$ | $\frac{R_{terMg}+C}{R_{tot}}$ |
| *Model 3* | $\frac{I+C}{R_{tot}}$ | $\frac{R_{terE}+C}{R_{tot}}$ | $\frac{C}{R_{tot}}$ | $\frac{R+R_{terE}+I+C}{R_{tot}}$ |
| *Model 4* | $\frac{I}{R_{tot}}$ | $\frac{R_{ter}+I}{R_{tot}}$ | $\frac{I}{R_{tot}}$ | $\frac{R_{ter}+I}{R_{tot}}$ |
| *Model 5* | $\frac{I+C}{R_{tot}}$ | $\frac{R_{terE}+C}{R_{tot}}$ | $\frac{C}{R_{tot}}$ | $\frac{R+R_{terE}+I+C}{R_{tot}}$ |
| *Model 6* | $\frac{I+C}{R_{tot}}$ | $\frac{R_{ter}+C}{R_{tot}}$ | $\frac{C}{R_{tot}}$ | $\frac{R+R_{ter}+I+C}{R_{tot}}$ |

**Table S3. Summary of bound and folded fractions for the models used in this study.** The table shows the expressions used to compute the *fraction bound* (corresponding to the total probability of bound RNA at a given SAM concentration ($S_{tot}$)), the *fraction folded*, the *fraction both bound* & *folded* and *fraction binding competent* for *Models 1-6*.
